# Characterization and evaluation of the anti-streptococcal activity of a novel zoocin A-like endopeptidase for potential veterinary applications

**DOI:** 10.64898/2026.10.23.760966

**Authors:** Marzena Nowacka, Magdalena Kaus-Drobek, Malgorzata Korzeniowska née Wiweger, Magdalena Gewartowska, Izabela Sabala

## Abstract

The increasing prevalence of antimicrobial resistance among streptococci necessitates the development of alternative antibacterial strategies against veterinary pathogens. Here, we report the first comprehensive characterization of ZooA_agal, a zoocin A-like bacteriolytic enzyme derived from *Streptococcus agalactiae* strain QMA0326. Both the proenzyme containing signal peptide (SP) pre-ZooA_agal and its mature variant containing the catalytic and target recognition domains (ZooA_agal) were successfully expressed and purified following the optimization of expression conditions. The enzyme displayed high selectivity toward bacteria possessing the A11.5 peptidoglycan chemotype (A1α-L-Lys-L-Ala_2_), which is common in streptococcal species, such as *S. dysgalactiae*, *S. agalactiae* or *S. uberis,* exhibiting a minimum bactericidal concentration of 0.5 μM and rapid dose-dependent lysis. Pre-ZooA_agal showed strong bacteriolytic activity across a broad pH range (6.0–11.0) and remained active in NaCl concentrations up to 500 mM. Its enzymatic activity was sustained in serum and milk and enhanced in the presence of Ca^2+^ and Mg^2+^ ions. Electron microscopy observations confirmed extensive structural damage to susceptible bacterial cells following enzyme treatment. Importantly, no toxicity of both pre-ZooA_agal and ZooA_agal was observed in vertebrate models. These findings highlight pre-ZooA_agal and ZooA_agal as a stable and highly selective enzybiotics with potential for the targeted control of streptococcal infections relevant to veterinary medicine.

## Introduction

Streptococci are key pathogens of both animals and humans, causing a broad spectrum of diseases. In livestock, species such as *Streptococcus agalactiae* and *Streptococcus uberis* are major contributors to bovine mastitis, resulting in substantial economic losses for the dairy industry worldwide^1–3^. In addition, species including *Streptococcus pyogenes* and *Streptococcus dysgalactiae* are responsible for numerous infections in both humans and animals, emphasizing the zoonotic potential and One Health relevance of this bacterial genus, where human, animal, and environmental health are closely interconnected^4,5^. In veterinary practice, β-lactam antibiotics such as penicillin G, amoxicillin, and ampicillin remain the first-line therapy for treating most streptococcal infections across various animal species. Nevertheless, increasing resistance to tetracyclines, macrolides, and lincosamides has been reported, particularly among swine-associated *Streptococcus suis* and mastitis-causing streptococci^6^. This growing prevalence of antimicrobial resistance emphasizes the urgent need for novel, targeted therapeutics suitable for veterinary applications.

Among the most promising alternatives to conventional antibiotics are enzybiotics, a class of peptidoglycan hydrolases (PGHs), enzymes that degrade the bacterial cell wall, ultimately resulting in cell lysis. Enzybiotics recognize specific cell wall components, bind to them and cleave bonds within the primary structural element of the wall – peptidoglycan^7,8^. They act upon contact, and their lytic effect is immediate. Unlike conventional antibiotics, PGHs typically exhibit narrow, species-level specificity, thereby minimizing the disruption of commensal microbiota while maintaining efficacy against multidrug-resistant pathogens and biofilms^9–11^. Furthermore, resistance to PGHs develops infrequently, and these enzymes have demonstrated favorable safety profiles for humans, animals, and the environment.

PGHs produced by bacteriophages (endolysins) or bacteria (autolysins and bacteriocins) have attracted considerable attention as next-generation antimicrobials. By cleaving specific bonds within the peptidoglycan layer, they rapidly lyse susceptible bacterial cells. Their targeted mode of action, combined with their efficacy against antibiotic-resistant bacteria, has stimulated extensive research into their therapeutic applications^12^.

Enzybiotics have already been evaluated across multiple therapeutic indications. Their efficacy has been demonstrated in animal models of infected wounds and mastitis^13^, as well as in clinical settings, including the topical treatment of atopic dermatitis^14,15^ (ClinicalTrial.gov identifier: NCT02840955), and, more recently, the decolonization of nasal *Staphylococcus aureus* carriage - including MRSA - with the phage lysin HY-133, currently under evaluation in a first-in-human Phase I trial^16^ (ClinicalTrial.gov identifier: NCT06290557). The significant progress made in the development and testing of enzybiotics in recent years highlights their strong potential as a modern and innovative alternative to conventional antibiotics.

Bacteriolytic enzymes targeting streptococci have also been reported. A prominent example is zoocin A, a streptococcolytic PGH originally identified in *Streptococcus equi* subsp*. zooepidemicus* strain 4881^17^. The enzyme displays species-dependent activity across streptococci, exhibiting particularly strong bactericidal effects against *S. pyogenes* and more moderate activity toward oral pathogens such as *Streptococcus mutans*^18^. Similar to many PGHs, mature zoocin A is organized into two functional domains: an N-terminal catalytic domain, which is responsible for peptidoglycan hydrolysis, and a C-terminal substrate-binding region, also referred to as the target recognition domain (TRD), which confers specificity toward susceptible streptococcal cell walls. These domains are connected by a flexible linker, which is rich in threonine and proline^19^.

Zoocin A functions as a D-alanyl-L-alanine endopeptidase, cleaving cross-links within the peptidoglycan network of the bacterial cell wall^20^. Its catalytic domain belongs to the M23 family of zinc-dependent endopeptidases and contains the characteristic HxxxD and HxH motifs that coordinate a zinc ion essential for enzymatic activity. Structural studies have confirmed that its M23 fold is similar to those found in Lss and LytM^21,22^. In contrast, the TRD of zoocin A represents a novel protein fold family with no known sequence homologs^23^. The structure of the isolated TRD was determined by NMR (PDB ID: 2LS0) and was shown to adopt a globular fold composed predominantly of antiparallel β-strands with a short C-terminal α-helix, forming a surface-exposed cleft that mediates binding to streptococcal peptidoglycan. The studies on recombinant TRD domain have confirmed that its peptidoglycan-binding takes place independently of the catalytic domain activity^19^. The TRD of zoocin A was also identified in the endolysin LyJH307, with an N-terminal NlpC/P60 catalytic domain. This enzyme exhibits rapid, dose-dependent lytic activity against *Streptococcus bovis* JB1, clearing cultures within minutes under acidic to neutral conditions (pH ∼5.5) and at rumen-relevant temperatures^24^. More recently, the zoocin A TRD domain was identified as the C-terminal target recognition domain of Lys572, a novel endolysin identified from the porcine intestinal virome. Lys572, comprising an N-terminal CHAP domain, a central Amidase-2 domain, and a C-terminal zoocin A TRD domain, exhibited potent activity against *S. suis* and *S. equi,* including effective removal of mature *S. suis* biofilms. Moreover, Lys572 retained activity across a broad range of pH and NaCl concentrations and reduced *S. suis* counts in refrigerated and thawed pork, highlighting the potential of TRD-containing enzymes as antibacterial agents for food biocontrol^25^.

Zoocin A is a zinc-dependent bacteriolytic enzyme and its activity is highly sensitive to the presence of metal ions. Chelation with 1,10-phenanthroline completely abolishes its activity, confirming the essential role of Zn^2+^ within the active site. In contrast, exposure to 5 mM concentrations of other divalent metal ions, such as Cu^2+^, Ca^2+^ and Mg^2+^, leads to a marked reduction in zoocin A activity. This inhibition is proposed to result from competition for metal-binding sites or conformational changes affecting catalysis rather than substrate recognition^18^.

Beyond the originally described *S. equi*-derived enzyme, the exploration of zoocin A homologs from other streptococcal species represents an important avenue for expanding the therapeutic arsenal of PGHs. Of particular veterinary relevance, several *S. agalactiae* strains isolated from bovine mastitis have demonstrated strong antagonistic activity against other streptococcal species, highlighting their competitive advantage within the mammary environment. Genomic analysis has revealed that these antagonistic strains produced multiple antimicrobial metabolites, including a zoocin A-like bacteriocin, which suggests a role for zoocin A homologs in inter-streptococcal inhibition^26^.

Despite this increasing understanding, several aspects of the mode of action of zoocin A remain incompletely defined. The site of hydrolysis within the peptidoglycan cross-bridge by zoocin A is established to be a bond between D-alanine residue of the stem peptide and the first L-alanine residue of the cross-bridge^20^, however some streptococci possessing the same PG chemotype are susceptible while others are not. This variation suggests that additional components of the bacterial cell wall may contribute to determining the specificity of, and/or regulating, zoocin A activity^27,28^.

In this study, we present the first comprehensive characterization of a novel zoocin A homolog derived from *S. agalactiae* strain QMA0326, examining both a truncated variant (ZooA_agal) and the full-length enzyme containing its native signal peptide (pre-ZooA_agal). We systematically evaluated the enzyme’s activity against bacterial strains possessing diverse peptidoglycan chemotypes, determined optimal pH and ionic conditions, and investigated its dependence on metal ion. Furthermore, we employed electron microscopy to visualize the cellular effects of enzyme treatment. Finally, we assessed the safety profile of ZooA_agal and demonstrated activity of the enzyme in the presence of serum and milk to evaluate its potential as a therapeutic agent for the targeted control of streptococcal infections, including bovine mastitis.

## Material and methods

### Identification of zoocin A homolog from Streptococcus agalactiae

The amino acid sequence of the LytM catalytic domain from *S. aureus* NCTC 8325-4 (residues 185–316; accession number O33599) was used as a query sequence for a BLAST (Basic Local Alignment Search Tool) search. This search was performed against streptococcal genomes within the non-redundant protein database available on the National Center for Biotechnology Information (NCBI) platform. BLAST parameters were set to default and sequences and values with significant similarity were selected for further analysis.

### Protein sequence analysis and structural prediction

The amino acid sequence of zoocin A homolog from *S. agalactiae* strain QMA0326 (GenBank accession: PWS69917.1) was retrieved from the NCBI database. This 299 amino acid protein is designated here pre-ZooA_agal.

Three-dimensional structural prediction was performed using ColabFold^29^, an online implementation of AlphaFold^30^. Default parameters were applied for structure prediction pipeline, which included multiple sequence alignment (MSA) generation using MMseqs2 and iterative refinement through recycling^31^. Five structural models were generated and the highest-ranked model was selected based on its predicted Local Distance Difference Test (pLDDT) score and overall confidence metrics. Structural visualization and analysis were conducted using PyMOL (PyMOL Molecular Graphics System, version 2.5.5, Schrödinger, LLC; http://pymol.org/).

### Bacterial strains used in the studies

The bacterial strains used in activity assays were purchased from: German Collection of Microorganisms and Cell Cultures (Leibniz Institute, DSMZ); Masaryk University, Czech Collection of Microorganisms (CCM); American Type Culture Collection (ATCC); and Polish Collection of Microorganisms (PMC) Institute of Immunology and Experimental Therapy PAN. The full list of bacterial strains with specified source is listed in the Supplementary Table S1.

Unless stated otherwise, the bacterial strains were cultivated in Tryptic Soy Broth (TSB) medium at 37°C with gentle shaking (at 90 rpm).

For cloning and protein overexpression, *Escherichia coli* BL21-Gold (DE3) strain (Agilent Technologies) was used in this study. For protein overexpression bacterial cells were grown in Lysogeny Broth (LB) Lennox formulation (Bioshop Canada Inc.).

### Plasmid construction and gene cloning

A synthetic gene encoding full length zoocin A from *S. agalactiae* (amino acids 1-299), codon-optimized for *E. coli* expression, was ordered from Eurofins Genomics. The Sequence and Ligation Independent Cloning (SLIC) method^32^ was used to generate pET28-6xHis-SUMO_pre-ZooA_agal expression construct for the production of full length zoocin A (pre-ZooA_agal) (with N-terminal 6xHis-SUMO tag). The truncated ZooA_agal construct (amino acids 31-299), was generated via PCR reaction using pET28-6xHis-SUMO_pre-ZooA_agal as a template effectively omitting the signal peptide sequence. The comprehensive list of the primers used in this study is provided in the Supplementary Table S2.

### Protein expression and purification

The sequence-verified plasmids were used to transform *E. coli* BL21-Gold (DE3) competent cells. The proteins were expressed overnight at 16°C after induction with 0.4 mM isopropyl β-D-1-thiogalactopyranoside (IPTG) when the cultures reached an OD_595_ of 0.6-0.8. Bacterial cells were resuspended in ice-cold buffer A (40 mM NaH_2_PO_4_ pH 7.0, 250 mM NaCl, 10% (v/v) glycerol, 20 mM imidazole). After 30-min incubation on ice the suspension was disrupted via sonication. The lysate was clarified by centrifugation at 20,000 × *g* for 40 min at 4°C and the resulting supernatant was loaded onto a column packed with cOmplete™ His-Tag Purification Resin (Roche) equilibrated with buffer A. The resin was sequentially washed with buffer A and then with the same buffer supplemented with 40 mM imidazole. Finally the target proteins were eluted using buffer A containing 300 mM imidazole. To remove the 6xHis-SUMO tag, the fusion proteins were digested overnight at 4°C by adding SUMO protease to desired fractions. The reaction mixture was applied to a size exclusion column (Hiload 16/600 Superdex 75 PG, Cytiva) equilibrated with a storage buffer (40 mM NaH_2_PO_4_ pH 7.0, 100 mM NaCl and 10% (v/v) glycerol). Protein purity was assessed after each purification step using Tricine-SDS PAGE gel electrophoresis. Pooled fractions of purified proteins were concentrated using a 10 kDa MWCO Centrifugal Filter Device (Sartorius), flash frozen in liquid nitrogen and kept at -80°C.

### Activity tests

#### Turbidity reduction assay (TRA)

A single colony of bacteria grown on a Tryptic Soy Agar (TSA) plate was inoculated into TSB and cultured overnight at 37°C with gentle shaking (90 rpm). The following day, fresh TSB medium was inoculated with 2% of the overnight culture and kept in a shaker at 37°C until bacterial culture reached an OD_595_ of 0.6-0.8. The cells were harvested at 4200 × *g* for 10 min at 4°C and resuspended in two distinct buffers - 50 mM glycine buffer (pH 8.0) either supplemented with or devoid of 100 mM NaCl - to the final cell density OD_595_ of 1.0, corresponding to around 1×10^8^ CFU/ml (Colony Forming Unit/ml). The lytic assays were performed in a sterile 96-well flat-bottom microplates for 1 h in that manner that 100 µl of each protein dilution was mixed with 100 µl of the prepared bacterial suspension, resulting in a total volume of 200 µl per well. The OD_595_ was monitored continuously every 2 min for 1 h with a brief 5 s orbital shaking step performed before each reading using a Bio-Rad microplate reader. Samples without enzyme added served as a control. The concentration of the enzyme used in experiments ranged from 0.5 to 2 μM. Each assay was performed in three technical and three independent biological replicates. A total of 22 bacterial strains (listed in the Supplementary Table S1) were screened to evaluate specificity of both pre-ZooA_agal and ZooA_agal using turbidity reduction assay (TRA). The lytic activity was calculated as the difference between the OD_595_ of the negative control and the OD_595_ of cells treated with the tested enzyme after a 1 h incubation period. The results are expressed as the percentage of OD_595_ reduction as compared to the control.

#### Effect of pH on lytic activity

To evaluate the optimal pH for enzymatic activity, turbidity reduction assays were performed across a pH range of 5 to 11. The assays were conducted at room temperature (RT) for 1 h using a final concentration of 1 µM against *S. agalactiae* cells suspended at approximately 1×10^8^ CFU/ml. To exclude the effect of ionic strength the set of buffers with different pH was prepared to match the conductivity of the reference 50 mM glycine buffer (pH 8.0). The following buffers were used: 10 mM sodium acetate pH 5.0, 2 mM sodium citrate (pH 6.0), 5 mM Tris-HCl (pH 7.0), 10 mM Tris-HCl (pH 8.0), 50 mM Tris-HCl (pH 9.0), 20 mM CAPS (pH 10.0) and 10 mM CAPS (pH 11.0).

#### Effect of ionic strength on enzymes activity

To investigate the effect of ionic strength on enzymatic activity, turbidity reduction assays were conducted in 50 mM glycine buffer (pH 8.0) supplemented with various NaCl concentration ranging from 0 to 1000 mM, while all other conditions remained identical as described in pH assay.

#### Drop killing assay

Bacterial cells were cultivated as for the turbidity reduction assay. The cell suspension was adjusted using McFarland Densitometer (Biosan) to 2.0 McFarland units (McF) which corresponds to approximately 2×10^8^ CFU/ml and further diluted in 50 mM glycine buffer pH 8.0 to reach cell density of around 1×10^6^ CFU/ml. The cells were mixed in a 1:1 (v/v) ratio with the enzyme and incubated at room temperature for 2 h. The final enzyme concentrations evaluated ranged from 0.5 to 2 μM. Following incubation, 10-fold serial dilutions were prepared in the same buffer and 5 μl of each sample was spotted onto TSA plates. The plates were incubated at 37°C overnight and the remaining viable colonies were enumerated the following day.

#### Effect of metal ions on lytic activity

To assess the effect of ions on enzymatic activity, reaction mixtures were prepared in a 96-well microplate. Each well contained 100 μl of bacterial suspension (1×10^8^ CFU/ml) of 4 strains*: S. dysgalactiae, S. agalactiae, S. pyogenes* and *S. canis,* and 100 μl of protein solution (2 μM), in the presence of 1 mM ions: either Ca^2+^ (CaCl_2_) or Mg^2+^ (MgCl_2_). Apart from the addition of ions, all reactions were prepared and monitored as described above in TRA section. Data analysis was carried out using GraphPad Prism software version 11.01.0 using two-way ANOVA with Dunnett’s post-hoc multiple comparisons test. Data is presented as a mean ± standard deviation. The differences were considered significant at P < 0.05.

#### Quantification of antibacterial activity via spot killing assay

Three mastitis-associated streptococcal strains (*S. agalactiae, S. dysgalactiae* and *S. uberis*) were tested to determine bacterial log_10_ reduction. Antibacterial activity of the enzyme was evaluated using a spot killing assay. Briefly, overnight cultures of the selected bacterial strains were diluted in sterile 50 mM glycine buffer pH 8.0 to the final cell density OD_595_ of 1.0, corresponding to around 1×10^8^ CFU/ml and further diluted to reach 1×10^7^ CFU/ml and 1×10^6^ CFU/ml. Bacterial suspensions were mixed at 1:1 ratio with the enzymes in the presence of 1 mM CaCl_2_ and incubated for 1 h at RT. Following incubation, samples were subjected to 10-fold serial dilutions in sterile glycine buffer to approximately 1×10^1^ CFU/ml. Five microliters (5 μl) of each sample were spotted onto TSA plates and incubated overnight at 37°C. Bacterial growth was assessed the following day, and the log_10_ reduction in viable counts was calculated relative to the untreated controls. The results were further expressed as the percentage reduction in viable bacterial counts.

#### Retention of antimicrobial activity under physiologically relevant conditions

Antimicrobial activity of the enzyme was evaluated against *S. agalactiae* under physiologically relevant conditions. Overnight bacterial cultures were diluted to a final cell density of OD_595_ of 1.0 (approximately 1×10^8^ CFU/ml) and further adjusted to reach 5×10^7^ CFU/ml. To assess the impact of biological environments, the enzyme was evaluated in fetal bovine serum (FBS), human serum, and ultra-high temperature (UHT) treated 0% fat milk at final concentrations of 100%, 50%, 25% and 12.5%, (v/v). Bacterial suspensions were mixed with the enzyme to obtain the final concentration of 5 µM in the presence of 1 mM CaCl₂ and incubated for 1 h and 24h at room temperature in the respective matrices. Following incubation, samples were subjected to 10-fold serial dilutions in sterile 50 mM glycine buffer (pH 8.0) to approximately 1×10^1^ CFU/ml. Five microliter (5 µL) aliquots were spotted onto TSA plates and incubated overnight at 37°C. Bacterial growth was assessed the following day, and the log reduction in viable counts was determined relative to the corresponding untreated controls. The results were further expressed as a log_10_ reduction corresponding to the percentage of bacterial eradication.

#### Determination of Minimum Bactericidal Concentration (MBC)

MBC determination was carried out using a spot killing assay. Three streptococcal strains (*S. agalactiae, S. uberis* and *S. dysgalactiae*) were cultured at 37°C to mid exponential growth phase and adjusted to an initial cell density of 1×10^5^ CFU/ml in sterile 50 mM glycine buffer (pH 8.0). The cells were mixed 1:1 with the enzymes of concentration ranging from 0.0011 to 1 µM and incubated for 24 h at RT. Suspensions were subjected to 10-fold serial dilutions in sterile glycine buffer to achieve a concentration of approximately 1×10^1^ CFU/ml. Subsequently, 5 μl aliquots of each dilution were spotted onto TSA plates and incubated overnight at 37°C. After incubation, the plates were examined for visible bacterial growth. MBC was defined as the lowest concentration of protein at which no growth was observed.

All activity assays were performed in both biological and technical triplicates.

### Electron microscopy

Scanning (SEM) and transmission (TEM) electron microscopy experiments were performed as previously described with slight modifications^33,34^. Briefly, *S. agalactiae* DSM6784 strain was cultured in TSB at 37°C with shaking at 80 rpm for 18 hours. The bacterial cells were then harvested by centrifugation and resuspended in a buffer containing 50 mM glycine buffer (pH 8.0) and 100 mM NaCl. The suspension was adjusted to a turbidity equivalent to 4 McFarland units (approximately 1×10^9^ CFU/ml). Cells were subsequently treated with ZooA_agal protein at the final concentrations of 100 nM or 200 nM for 1 h at room temperature. Untreated bacterial suspensions in the same buffer were used as a control.

For SEM analysis, negative control as well as bacterial cell suspensions treated with ZooA_agal were loaded onto 0.4 µm-pore size membrane filters (Isopore, 13 mm diameter). Bacteria were fixed in 2.5 % glutaraldehyde in 0.1 M cacodylate buffer for 2 h, followed by washing in the same buffer. Samples were then fixed in 0.2% osmium tetroxide and dehydrated in an ethanol gradient. Subsequently samples were dried using liquid CO₂ in a critical point dryer (B7020, Agar Scientific) and spray-coated with a thin layer of gold using a sputter coater (SC7620, POLARON).

The images presented herein were acquired with a JSM-6390LV scanning electron microscope (Jeol, Tokyo, Japan).

For TEM, the cell pellets were treated and fixed as for SEM, but were kept in solution instead on the filters. The dehydrated bacteria pellets were embedded in Embed 812 epoxy resin (Serva, Heidelberg, Germany). Ultrathin sections were stained for 15 minutes with 2% uranyl acetate and 5 minutes in lead citrate. Imaging was performed using a JEM-1011 EX transmission electron microscope (Jeol, Tokyo, Japan) equipped with a MORADA camera and iTEM v.1233 image analysis software (Olympus Soft Imaging Solutions, Münster, Germany).

### In vitro cytotoxicity assays

Mouse fibroblasts (NIH 3T3 cell line) and human keratinocytes (HaCaT cell line) were cultured in Dulbecco’s Modified Eagle Medium (DMEM) supplemented with 10% fetal bovine serum (FBS) and 1% penicillin-streptomycin. Cells were kept in ø 90 mm Petri dishes at 37°C in a humidified atmosphere containing 5% CO_2_ and subcultured two to three times a week. At approximately 80% confluence, cells were washed with 10 ml phosphate-buffered saline (PBS), and detached using 1 ml of TrypLE enzyme solution (Gibco).

For the cytotoxicity assay, phenol-red-free DMEM medium supplemented with 10% FBS was used to prepare the cell suspension, and adjust its density to 1×10^5^ cells/ml. 100 µl aliquots of the cell suspension were seeded into wells in columns 1-6 of a 96-well plate. For background control, columns 7-12 were filled with 100 µl of medium without cells. After 24 h of incubation, the culture medium was gently replaced with freshly prepared and filtered through a 0.22 µm filter: DMEM with 10% FBS (positive control), DMEM with 10% FBS and 10% ethanol (negative control), or DMEM with 10% FBS and ZooA_agal (treatment). Cells were further incubated at 37°C in 5% CO_2_ for an additional 22-24 h. Subsequently, 10 μl of MTT dye (5 mg/ml stock in PBS) was added to a final concentration of 0.5 mg/ml, and plates were incubated for an additional 2-3 h to let the formazan crystals form. To dissolve the crystals and read the absorbance of formazan, 100 µl of dimethyl sulfoxide (DMSO) was added to each well, and plates were gently agitated for 15-20 min. Absorbance was measured at 570 nm using a spectrophotometer. Cell viability was expressed as a percentage relative to the untreated positive control after subtraction of background absorbance. The experiment was conducted in three independent biological and six technical replicates per condition, using cells at early passages.

### In vivo toxicity tests

The greater wax moth (*Galleria mellonella*) larvae were obtained from a local vendor. Healthy-looking larvae at the final instar stage (approximately 2 cm in length) were selected to be maintained in ø 90 mm Petri dishes at room temperature in the dark without food for 24 h prior experiment. Groups of 10 larvae were injected in the abdomen adjacent to the last proleg using 0.5 ml insulin syringe with a 30-gauge needle. Each larva received 10 µl of either buffer (control) or ZooA_agal solution (1 or 4 µM in buffer, treatment). After injection larvae were incubated at 28°C in a humidified environment in the dark for one week. Larvae were inspected daily for melanisation, developmental progression, and survival. At the end of the experiment, survival was assessed based on the presence of viable larvae, cocoons, and pupae.

Adult zebrafish (*Danio rerio*) AB^OMD^ line purchased from Experimental Medicine Center (OMD) Lublin were housed and bred in the zebrafish facility at Mossakowski Medical Research Institute (MMRI), Polish Academy of Sciences as described in Kaus-Drobek et al., (2025)^34^. For the experiment, embryos originating from spontaneous crosses were collected, rinsed with E3 medium (5 mM NaCl, 0.17 mM KCl, 0.33 mM CaCl2, 0.33 mM MgSO4, pH 7.2^35^) and subjected to a morphological quality assessment. Only clutches containing >90% fertilized, normally developing embryos were used. Groups of 12 embryos at < 4 h post fertilization (hpf) were transferred to ø 30 mm Petri dish containing 2 ml of either E3 medium (untreated control) or 4 µM ZooA_agal in E3 medium (treatment). Embryos were incubated at 28°C for four days. Development was monitored daily under a stereomicroscope, and the following parameters were scored: survival rate, coagulation of embryos, absence of somite formation, lack of tail-bud detachment from the yolk sac, absence of a heartbeat, and other developmental abnormalities. Any dead embryos were removed, and medium was replaced daily with fresh E3 medium either supplemented with or devoid of ZooA_agal.

## Results

### Basic characterisation of the protein

A BLAST search using the catalytic domain sequence of LytM from *S. aureus* NCTC 8325-4 (residues 185-316; accession number O33599) resulted in identification of a 299 aa long putative zoocin A homolog in *S. agalactiae* QMA0326 (GenBank: PWS69917.1). The protein was named pre-ZooA_agal.

InterProScan (EMBL-EBI^36^) analysis of the protein sequence identified an N-teminal signal peptide (residues 1–27), an M23 peptidase catalytic domain (residues 49–139), and a target recognition domain (TRD) (residues 194-299). These functional domains are connected via a threonine and proline–rich linker which likely imparts flexibility between the domains, thereby ensuring the proper orientation of the recognition domain for substrate binding while allowing catalytic access to the peptidoglycan network^17^ (Fig. 1A).

**Figure 1.**
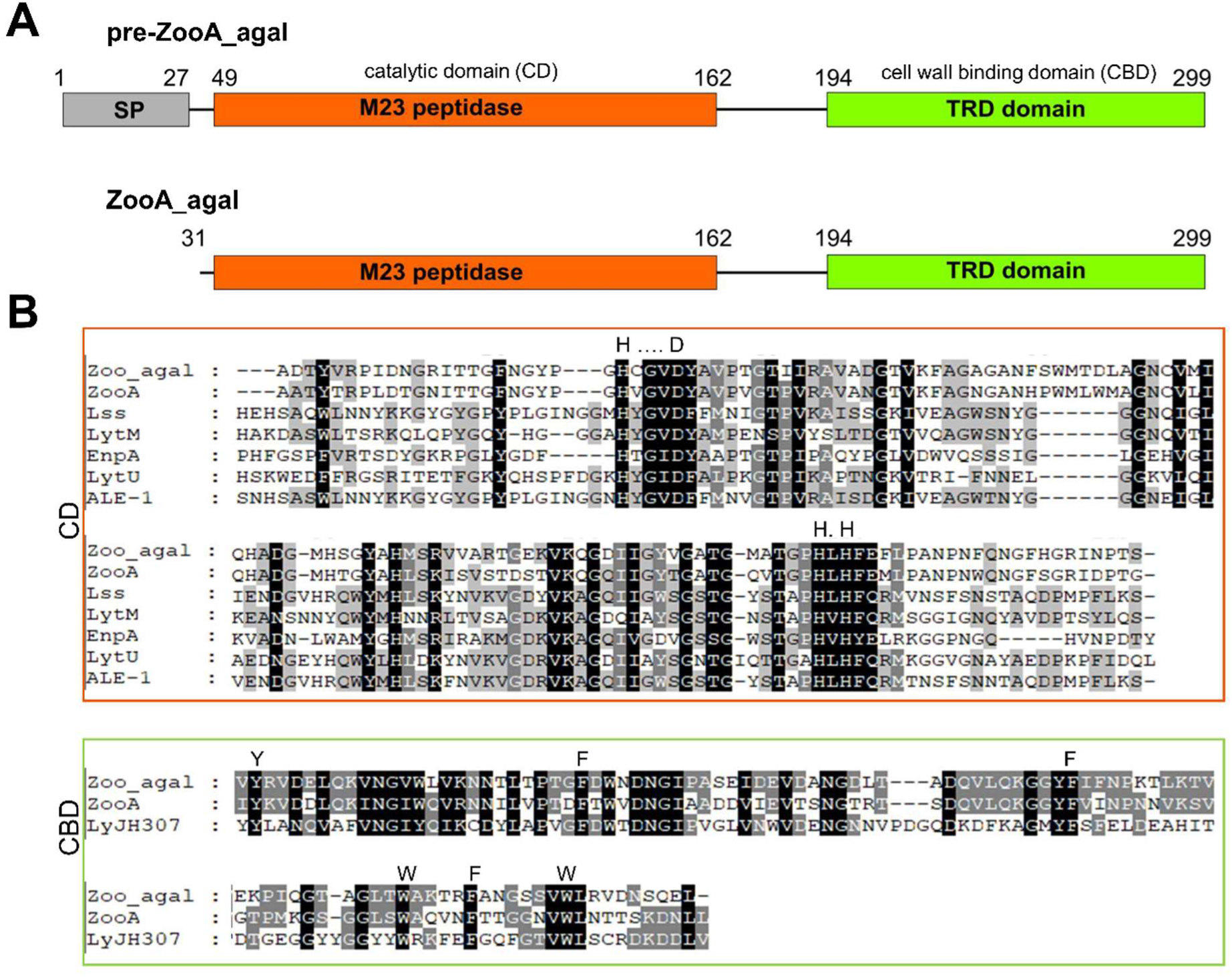
A) Domain architecture of the native pre-ZooA_agal and ZooA_agal constructs; SP-signal peptide. B) Sequence conservation in M23 peptidases; CD-catalytic domain; CBD-cell wall binding domain. Zinc-binding motifs HxxxD / HxH are indicated above the CD sequences; residues involved in the hydrophobic groove formation are indicated above CBD sequences.

Comparative analysis of pre-ZooA_agal with other well characterized M23 peptidases - including zoocin A, lysostaphin (Lss), LytM, EnpA, LytU and ALE-1, demonstrated high conservation of the characteristic HxxxD and HxH zinc binding motifs (Fig. 1B). Furthermore, sequence comparison of TRD domain of pre-ZooA_agal with TRDs of zoocin A from *Streptococcus equi* and LyJH307 from *Streptococcus equinus* enabled the identification of conserved residues contributing to the formation of a hydrophobic binding groove, a structural feature preserved across TRD domains^23^ (Fig. 1B).

Based on this data and to assess the potential role of the signal peptide in modulating the protein’s enzymatic activity as previously reported^37^, two expression constructs were designed: pre-ZooA_agal - expressing full length zoocin A (aa 1-299) and its truncated version lacking the signal peptide - ZooA_agal (aa 31-299).

Due to the initial difficulties in overexpression and solubility of the desired protein, a series of rational and empirical modifications were implemented to improve the yield of protein production and the subsequent handling without compromising its activity. Among others, N-terminal 6xHis_SUMO fusions were introduced reducing protein aggregation during the purification process.

### Predicted structural features of pre-ZooA_agal

The predicted AlphaFold model of the pre-ZooA_agal three-dimensional structure adopted well-defined and compact fold with overall high model confidence. The majority of residues exhibited high pLDDT values (predicted Local Distance Difference Test), indicating reliable structural prediction across most of the protein sequence. The N-terminal catalytic domain (M23 peptidase) is predominantly composed of antiparallel β-sheets and a single ɑ-helix. Six anti-parallel β-strands and four loops form a binding groove (Supplementary Fig. S1). The C-terminal region of the protein comprises the Target Recognition Domain (TRD) composed of six anti-parallel β-strands and a single ɑ-helix. The two domains are connected by a flexible 32-amino-acid linker.

Lower confidence regions were primarily located at the N-terminal segment (signal peptide), the linker region between M23 and TRD domains, and several surface-exposed loop regions, which may indicate structural flexibility or intrinsic disorder in these areas. Overall, the predicted model suggests a stable globular protein structure, with no large disordered domains detected.

### Specificity of the proteins

The lytic activity of pre-ZooA_agal and ZooA_agal proteins was initially assessed using a turbidity reduction assay (TRA) against a panel of 22 bacterial strains encompassing diverse cell wall types and peptidoglycan compositions (Supplementary Table S1).

Significant bacteriolytic activity, defined as an optical density reduction of at least 25% relative to the untreated control, was observed for eight of the tested bacterial species (Fig. 2). All susceptible strains were Gram-positive bacteria possessing either the A11.5 (A1α-L-Lys-L-Ala_2_) or A11.24 (L-Lys-L-Thr-Gly) peptidoglycan chemotype according to BacDive database^38^. Both enzymes were the most effective in lysing *Enterococcus faecalis* and several streptococcal species with alanines in their cross bridges, namely *S. dysgalactiae*, *S. agalactiae, S. salivarius* and *S. pyogenes* reducing the OD_595_ by at least 50%. A lower efficacy was observed against *S. uberis* suggesting that factors beyond the type of cross-bridge may affect enzymes efficacy. In general, the bacteriolytic efficacy of both enzymes was comparable, with the notable exception of *S. canis* having threonine and glycine in the cross-bridge, that was more sensitive to pre-ZooA_agal treatment.

**Figure 2.**
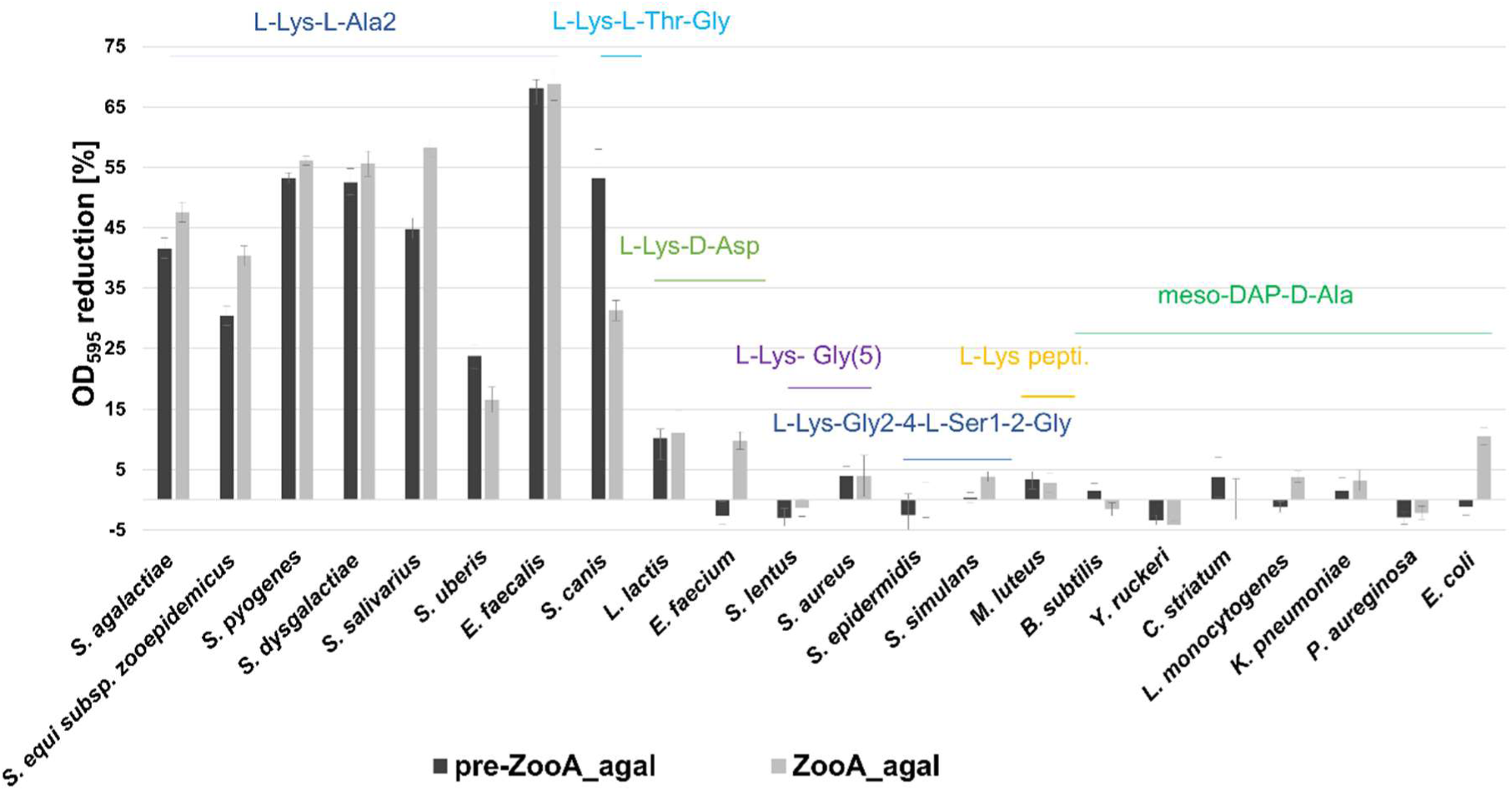
Lytic activity of pre-ZooA_agal and ZooA_agal against selected bacterial strains with the various types of peptidoglycan. Lytic activity, determined in TRA assay, is expressed as the percentage of reduction in OD_595_ relative to the control (without enzyme) after 1 h incubation with 1 µM protein at room temperature in 50 mM glycine buffer (pH 8.0) with 100 mM NaCl. A comprehensive list of the tested strains and their respective cell wall characteristics is provided in Supplementary Table S1. The sequences of peptidoglycan crossbridges are indicated above the corresponding bars.

Importantly, enzymes treatment did not cause any significant reduction in optical density for either the Gram-negative bacteria or the Gram-positive strains characterized by a direct cross-bridge or cross-bridges containing Gly, Gly-Ser, or Asp.

### Determination of optimal conditions for antibacterial activity of pre-ZooA_agal and ZooA_agal

The bacteriolytic activity of both variants, pre-ZooA_agal and ZooA_agal, was evaluated across a broad range of pH and ionic strength conditions. Turbidity reduction assays were conducted using 1 µM enzymes incubated with *Streptococcus agalactiae* DSM 6784 (OD_595_ = 1; ∼1×10^8^CFU/ml). To evaluate pH dependence, low ionic strength buffers of pH ranging from 5.0 to 11.0 were used. As shown in Fig. 3A, the optimal conditions for pre-ZooA_agal variant was between pH 7.0 and 9.0, whereas the optimum for the ZooA_agal variant it was shifted towards lower pH (6.0 – 8.0). Furthermore, while ZooA_agal demonstrated high sensitivity to alkaline environments, exhibiting a significant drop in the activity starting at pH 9.0, the pre-ZooA_agal variant sustained lytic activity up to pH 11.0.

**Figure 3.**
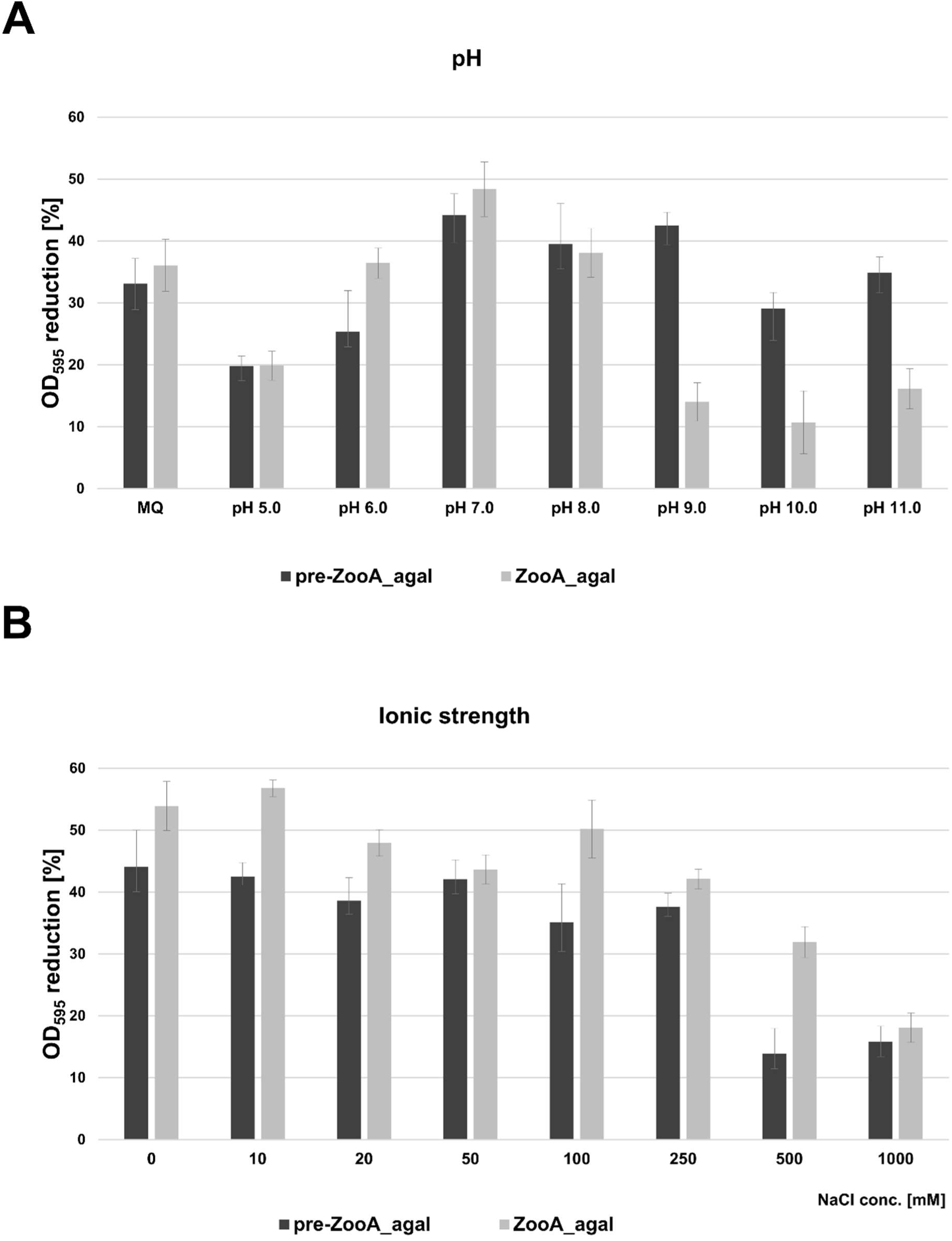
The effect of pH and ionic strength of buffers on the lytic activity of pre-ZooA_agal and ZooA_agal. The lytic performance against *S. agalactiae* DSM 6784 was determined via TRA assay after 1h incubation at room temperature. A) Activity assessed across varying pH (exact conditions are detailed in Materials and Methods section) and in Milli-Q water (MQ). B) Activity of 1 µM enzymes evaluated in 50 mM glycine buffer (pH 8.0) in the presence of increasing NaCl concentration.

The effect of ionic strength on pre-ZooA_agal and ZooA_agal activity was investigated in 50 mM glycine buffer (pH 8.0) supplemented with varying concentrations of NaCl ranging from 0 to 1000 mM (Fig. 3B). Both enzyme variants retained high activity in the presence of up to 250 mM of NaCl. Notably, in the presence of 500 mM NaCl ZooA_agal remained highly active, whereas the lytic efficacy of pre-ZooA_agal was strongly attenuated. In the presence of 1000 mM NaCl activity of both variants was markedly reduced.

The influence of metal ions on enzymatic activity was assessed via turbidity reduction assay in the presence of Ca^2+^ or Mg^2+^ ions (Fig. 4). The procedure was performed on four bacterial strains: *S. dysgalactiae* DSM 20662, *S. pyogenes* DSM 7418, *S. agalactiae* DSM 6784 and *S. canis* DSM 7148. The lytic activity of pre-ZooA_agal variant was not affected by the presence of divalent metal ions. The addition of CaCl_2_ and MgCl_2_ supported the highest levels of enzymatic activity of ZooA_agal variant, particularly in the case of *S. canis*, that was least susceptible among the tested strains in the control conditions (Fig.4).

**Figure 4.**
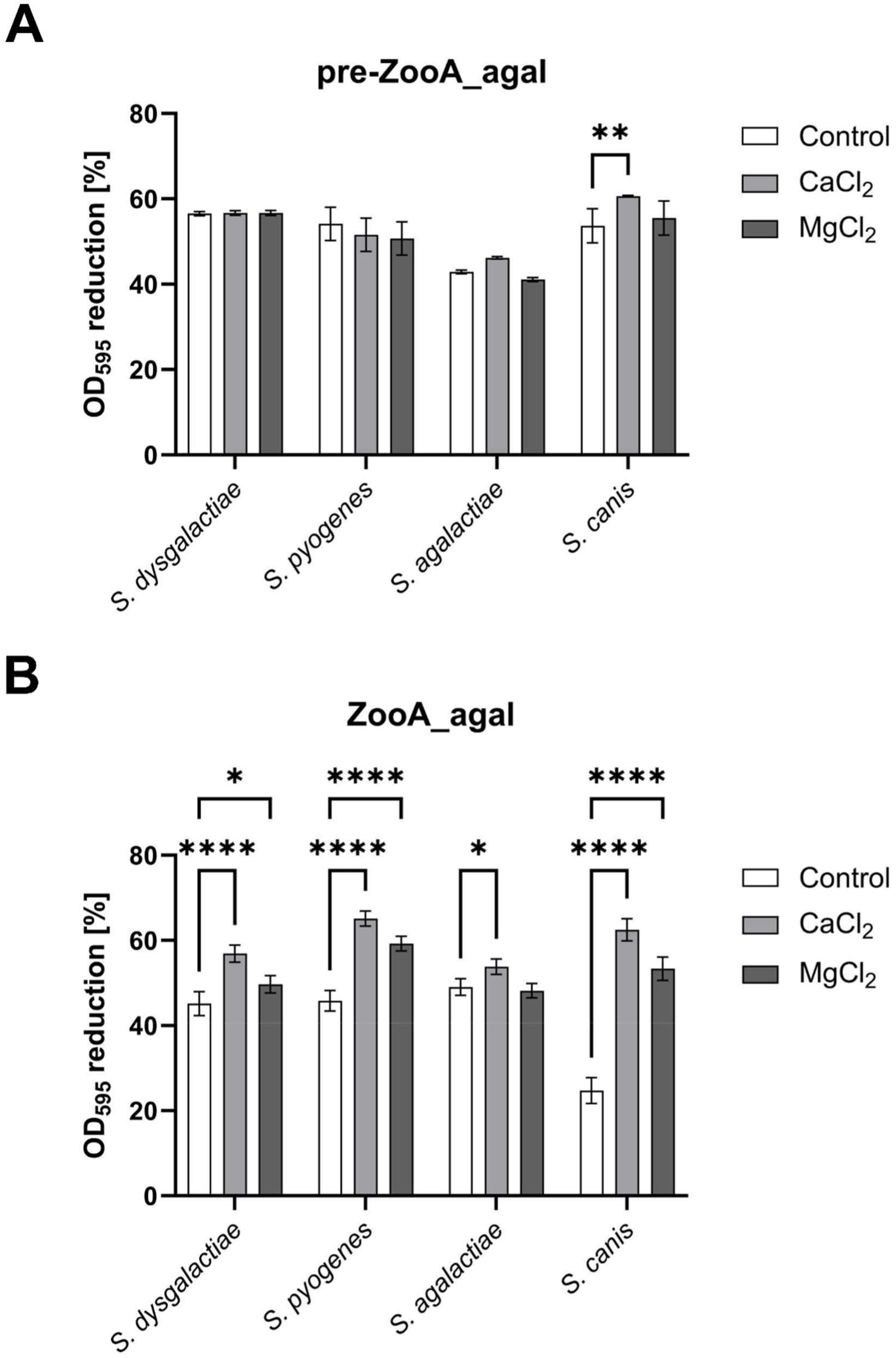
Metal ion dependence of pre-ZooA_agal and ZooA_agal activity. The lytic performance was determined against the following strains: *S. dysgalactiae* (orange), *S. pyogenes* (green), *S. agalactiae* (grey) and *S. canis* (yellow) via the turbidity reduction assay in 50 mM glycine buffer (pH 8.0) (control) or supplemented with a 1 mM final concentration of each specific metal ion. Bacterial cells were treated with 1 µM enzyme and OD_595_ was monitored for 1 h at room temperature. Two-way ANOVA with Dunnett’s post-hoc multiple comparisons test was carried out using GraphPad Prism software and is presented as a mean ± standard deviation. Statistical significance is depicted as **** - P < 0.0001, *** - P < 0.001, ** - P < 0.01, * - P < 0.05.

### Antibacterial efficacy against mastitis-associated Streptococcus species

To directly demonstrate the enzyme efficacy in eradication of mastitis related bacteria, the ZooA_agal variant was tested against *S. agalactiae, S. dysgalactiae,* and *S. uberis* using a spot killing assay. Bacterial cells at initial concentrations of 1×10^6^ CFU/ml or 1×10^7^ CFU/ml were treated with 1µM enzyme in 50 mM glycine buffer (pH 8.0) supplemented with 1mM CaCl_2_ for 1 h at room temperature prior to serial dilution and spotting onto TSA plates. Enzyme treatment resulted in a significant reduction in viable bacterial counts across all tested strains compared with untreated controls (Fig. 5). The efficacy of the bacterial reduction was species-dependent, with *S. agalactiae* being the most susceptible, demonstrating a 4 log_10_ reduction in viable counts, corresponding to 99.99% bacteria eradication. In contrast, *S. uberis* showed lower sensitivity to enzymatic treatment, yielding a one log_10_ reduction corresponding to eradication of ca. 90% of bacterial cells. *S. dysgalactiae* exhibited moderate susceptibility, with a 2 log_10_ reduction corresponding to the elimination of 99% of bacterial cells. These results demonstrate the robust and consistent efficacy of the enzyme against major mastitis-associated streptococci.

**Figure 5.**
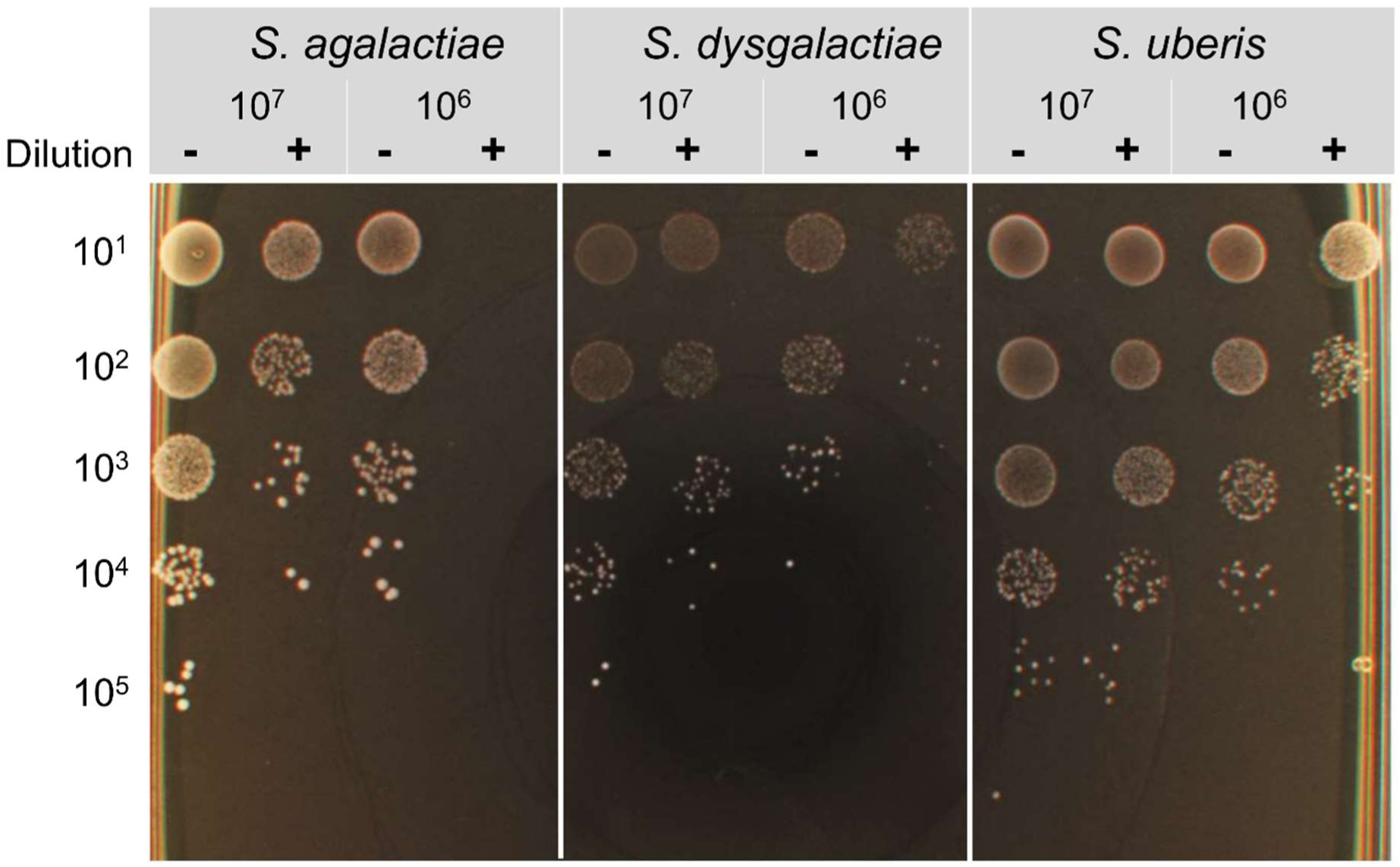
Antibacterial efficacy of ZooA_agal against mastitis-associated *Streptococcus* species. The bactericidal activity of the ZooA_agal against *S. agalactiae, S. dysgalactiae,* and *S. uberis* was determined by spot killing assay. Bacterial suspensions at initial concentrations of 1×10^6^ CFU/ml or 1×10^7^ CFU/ml were incubated with 1 µM enzyme in 50 mM glycine buffer (pH 8.0) supplemented with 1 mM CaCl_2_ for 1 h at room temperature prior to serial dilution and spotting on TSA plates.

### Retention of antimicrobial enzyme activity under physiologically relevant conditions

As it is known that activity of bacteriolytic enzymes can be substantially inhibited by serum or milk, we have first tested efficacy of the enzymes in the presence of human serum, bovine serum or milk. We observed a very significant, three or four logs reduction of CFU/ml after just 1 h treatment but only in the presence of 12.5% or 25% of tested matrices, while in 50% or full matrices the activity was not detectable (Supplementary Fig. S2A). Therefore, the treatment was extended to 24 h, to observe the antibacterial activity of the enzyme with measurable reductions in viable bacterial counts (Supplementary Fig. S2B).

The antibacterial effects were detected in milk with 3-4 log_10_ reductions in viable bacterial counts corresponding to eradication of 99.9% - 99.99% of bacterial cells. These findings demonstrate that the enzyme remains active under physiologically relevant conditions, supporting its potential applicability in environments that more closely mimic *in vivo* conditions.

### Minimal Bactericidal Concentration (MBC)

The minimum bactericidal concentration (MBC) of the ZooA_agal was determined using a spot killing assay. *S. agalactiae, S. uberis* and *S. dysgalactiae* cells were treated with serial dilutions of the enzyme ranging from 1 to 0.0011 µM, and the bactericidal efficacy was assessed after 24 h incubation at room temperature. The final MBC value, defined as the lowest protein concentration that totally prevented visible microbial growth, was determined to be 0.5 µM for all three tested strains, while clear inhibition was observed at concentrations as low as 0.125 µM (Supplementary Fig. S3).

### Ultrastructural analysis via electron microscopy

Electron microscopy showed that the untreated *S. agalactiae* cells retained normal morphology with intact cell walls and homogeneous cytoplasmic contents (Fig. 6A, B, E, F). In contrast, cells incubated with ZooA_agal exhibited profound ultrastructural alterations. Scanning electron microscopy (SEM) images revealed clear evidence of cytoplasmic leakage and accumulation of cellular debris in the extracellular space (Fig. 6C, D). Transmission electron microscopy (TEM) further confirmed cytoplasmic leakage and showed extensive damage to the bacterial cell walls, including visible ruptures (Fig. 6G–J). In addition, numerous electron-lucent cells with disrupted and disorganized cytoplasmic material were detected (Fig. 6G–J), together with remnants of fragmented bacterial cell walls (Fig. 6J).

**Figure 6.**
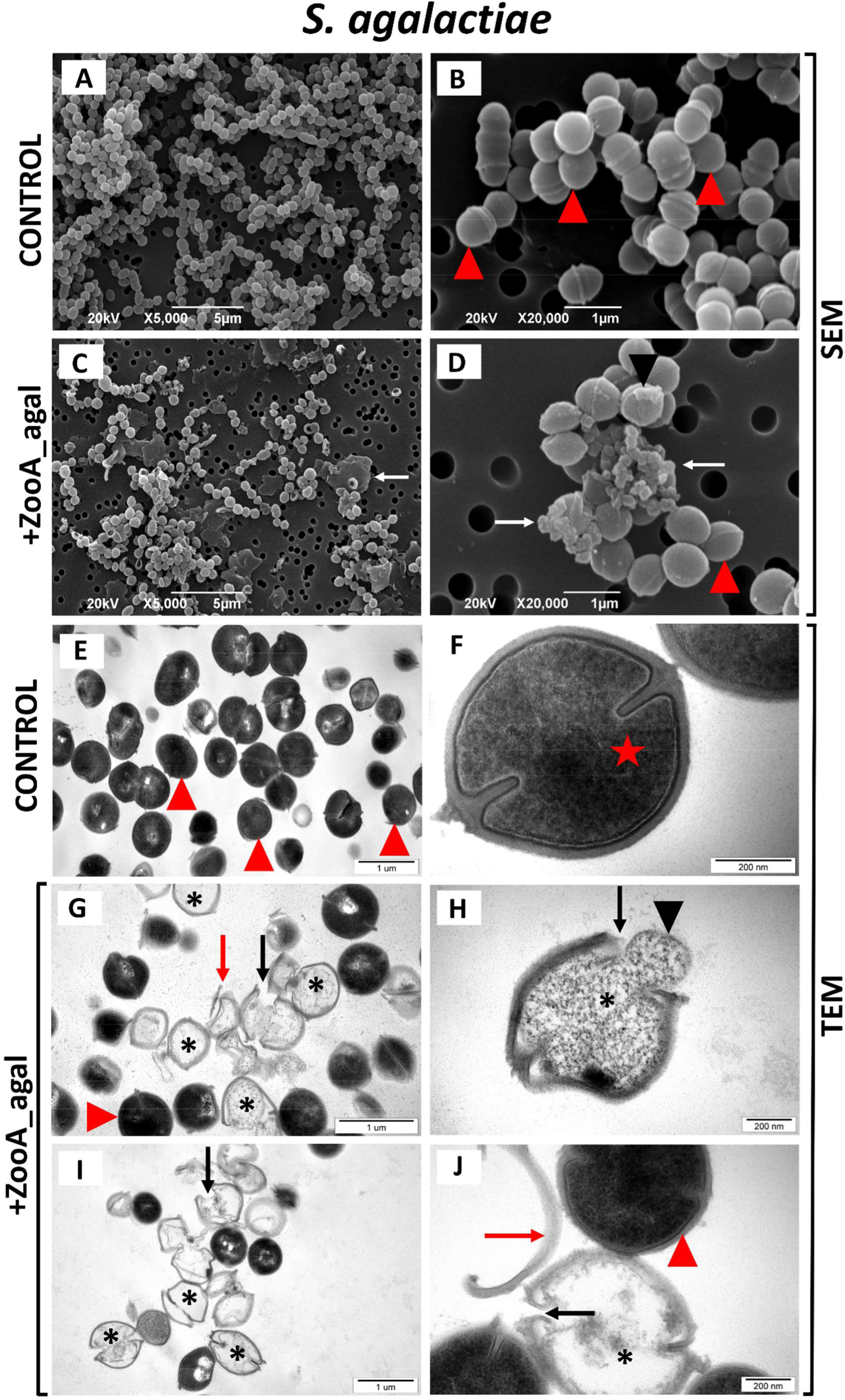
Ultrastructural analysis of *S. agalactiae* cells following treatment with ZooA_agal. Scanning electron microscopy (SEM) and transmission electron microscopy (TEM) images of *S. agalactiae* are presented. A, B - SEM images of untreated control cells. C, D - SEM images of cells incubated with ZooA_agal. E, F - TEM images of control cells. G–J - TEM images of cells incubated with ZooA_agal. Red arrowheads indicate ultrastructurally intact cells; white arrow - cellular debris; red star - an intact dividing cell; black arrowheads - leakage of intracellular contents; black arrows - ruptures in the bacterial cell wall; red arrows - remnants of the cell wall; asterisks - electron-lucent cells with disrupted cytoplasmic contents.

Taken together, the ultrastructural perturbations observed upon ZooA_agal exposure strongly demonstrate that this enzyme destroys bacterial cell walls, resulting in membrane disintegration and disruption of intracellular organization, thereby promoting cytoplasmic efflux and impairing bacterial viability.

### In vitro and in vivo toxicity assessment of pre-ZooA_agal and ZooA_agal

The cytotoxicity of pre-ZooA_agal and ZooA_agal was evaluated *in vitro* using the MTT assay on NIH 3T3 mouse fibroblasts and HaCaT human keratinocytes (Fig. 7A). Cells cultured in DMEM served as untreated controls, and all viability values were expressed as a percentage relative to this control. Exposure to 10% ethanol resulted in a marked reduction in cell viability in both cell lines, with viability decreasing to approximately 30% for 3T3 and 45% for HaCaT cells, confirming the expected cytotoxic effect of ethanol. In contrast, treatment with pre-ZooA_agal at concentrations of 1 µM and 4 µM did not significantly affect cell viability in either cell line, with values remaining comparable to the untreated control group. Similarly, exposure to 1 µM or 4 µM ZooA_agal maintained or slightly increased cell viability in both NIH 3T3 and HaCaT lines. These findings indicate that neither pre-ZooA_agal nor ZooA_agal exert cytotoxic effects on mammalian cells at the tested concentrations.

**Figure 7.**
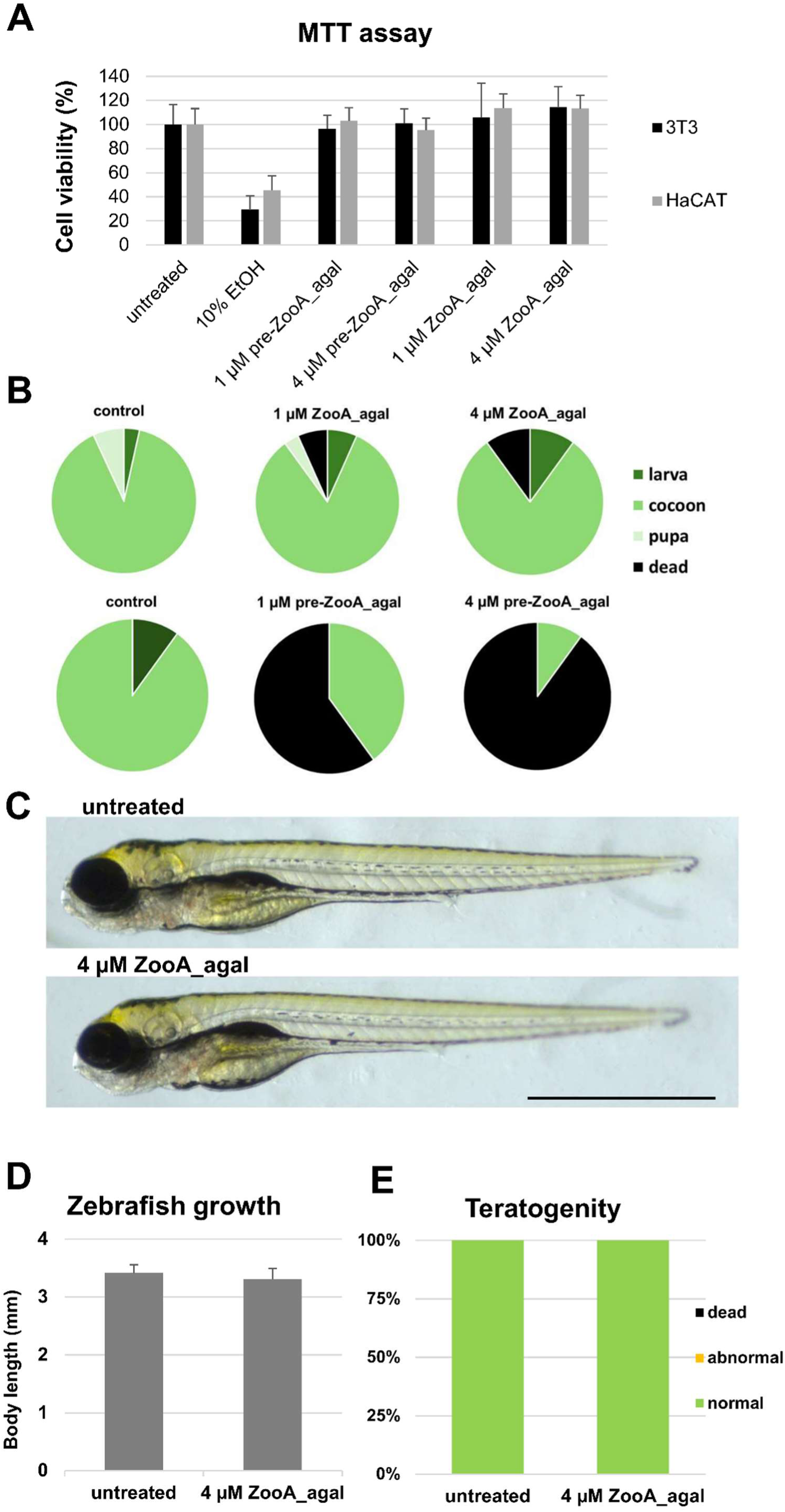
*In vitro* and *in vivo* toxicity evaluation of ZooA_agal and pre-ZooA_agal. A) MTT cell viability assay in which 3T3 mouse fibroblasts and HaCaT human keratinocytes were treated with ZooA_agal or pre-ZooA_agal at concentrations of 1 µM and 4 µM for 24 h. Cells cultured in DMEM served as untreated controls (100% viability), and viability values were expressed as percentages relative to this control. Exposure to 10% ethanol (EtOH) was used as a positive cytotoxic control. Data represent mean ± standard deviation from three independent experiments, each in 3-9 technical repetitions. B) Toxicity assay on the great wax moth larvae. Pie charts represent the proportions of phenotypes after 7 days post injection with 0 µM (mock injected control), 1 µM, and 4 µM ZooA_agal or pre-ZooA_agal.. Categories include larva (dark green), cocoon (light green), pupa (pale green), and dead (black). Higher concentrations of pre-ZooA_agal increased mortality and reduced cocoon formation, demonstrating a dose-dependent toxic effect. The experiment on *G. mellonella* was repeated two-to-three times with similar results, n=10. C-E) An example of acute toxicity assay on zebrafish embryos. C) Unchanged morphology of zebrafish larvae exposed to 4 µM ZooA_agal for four days at 28°C. photos taken under Stemi508 Zeiss stereomicroscope present lateral views larvae at 96 hpf. D) Bar plot demonstrates unchanged body length of zebrafish at 96 hpf indicative of ZooA_agal having no effect on zebrafish embryo development. E) Distribution of developmental scores, showing unaffected development and survival, suggesting non teratogenic effect of ZooA_agal on zebrafish embryos. The experiment on zebrafish was repeated three times with similar results. Statistical significance was determined using an unpaired Student’s t-test. n=12, scale bar=1 mm.

Injection with pre-ZooA_agal was highly toxic to *G. mellonella* larvae in a clear dose-dependent manner, causing 50% mortality at 1 µM and reaching 90% mortality at 4 µM concentration. By comparison, ZooA_agal induced only a modest increase in mortality, which remained below 10% (Fig. 7B) Surviving larvae developed normally, and the minor differences observed in the proportions of free larvae, cocoons, and pupae, at the end of the experiment, likely reflected subtle variations in the initial developmental stage rather than treatment-induced effects. Due to the pronounced observed toxicity of pre-ZooA_agal, only ZooA_agal was selected for subsequent tests on zebrafish embryos.

Zebrafish embryos exposed to ZooA_agal, as well as untreated controls, survived throughout the monitoring period with no observable mortality (Fig. 7C-E). At 4 days post-fertilization (dpf), larvae from both groups had similar body length, demonstrating that exposure to 4 µM ZooA_agal did not impair growth or developmental progression. Throughout the 4-day observation period, no adverse phenotypes mentioned in the OECD Test 236 for Fish Embryo Acute Toxicity (FET) test guidelines - embryo coagulation, defective somite formation, failure of tail-bud detachment from the yolk sac, absence of heartbeat, or any other developmental abnormalities were observed in either the control or in the treatment group. Collectively, these *in vitro* and *in vivo* results indicate that ZooA_agal variant does not induce acute embryotoxic or teratogenic effects under the conditions tested.

## Discussion

### Streptococci in mastitis, antimicrobial resistance, and enzybiotics as alternative therapies

Intramammary antibiotic administration remains the conventional strategy for clearing bacterial infections responsible for bovine mastitis. However, the extensive and widespread use of these drugs has accelerated a global resistance crisis^39^. Streptococcal strains, which represent major etiological agents of mastitis, exhibit increasing resistance to standard veterinary treatments. Furthermore, these bacteria can form resilient biofilms within the mammary gland that act as physical shields, blocking traditional drugs from reaching their targets. Moreover, *S. uberis* can internalize within bovine cells to evade immune responses and treatment^40^, frequently leading to chronic, recurrent infections.

To address this therapeutic challenge, current research is increasingly focused on non-antibiotic alternatives, such as bacteriophage-derived endolysins and bacterial autolysins^41^. These enzymes target and cleave specific bonds within the peptidoglycan of bacterial cell walls. Although pioneering studies evaluated wild-type enzymes like PlyC and Ply700^42,43^, their lytic activity is often attenuated by the complex composition of milk which contains inhibitory lipids and protein and high concentrations of various ions, like calcium or magnesium. To overcome these matrix-induced limitations, current research focuses on engineered chimeric lysins with enhanced lytic efficacy and stability in physiological environments^41^.

### Pre-ZooA_agal and ZooA_agal as targeted alternatives to conventional antibiotics

This study provides a comprehensive characterization of pre-ZooA_agal, the zoocin A homolog from *Streptococcus agalactiae* that effectively targets *S. agalactiae*, *S. uberis*, *S. dysgalactiae* and *E. faecalis*, thereby directly tackling the primary etiological agents responsible for bovine mastitis resulting in severe economic losses in the dairy industry. Deletion of the pre-ZooA_agal signal peptide yielded a biochemically robust, non-toxic variant (ZooA_agal). Both enzyme variants demonstrate potent lytic activity across a broad pH range (6.0-11.0 for pre-ZooA_agal, 6.0 – 8.0 for ZooA_agal) and under a wide spectrum of ionic conditions (up to 500 mM NaCl). Their low minimal bactericidal concentration (MBC) values coupled with sustained activity in the presence of calcium ions, milk and serum position these enzybiotics as promising alternative for conventional antimicrobial therapies in veterinary and human medicine.

### Specificity

To determine the specificity and lytic profile of the enzymes we assessed their activity against a diverse panel of 22 bacterial strains characterized by distinct cell wall architecture and peptidoglycan chemotypes, which is routinely employed in the screening workflows of our laboratory, thereby ensuring consistency that supports reproducibility and enables reliable comparative analyses^34,44^. Importantly, this panel includes veterinary pathogens responsible for animal infections, primarily bovine mastitis isolated from milk or infected udders, among others staphylococci (e.g. *S. aureus*) and streptococci (e.g *S. uberis, S. galactiae, S. disgalactiae*). Additionally, the panel includes strains isolated from livestock products and their veterinary environments, such as *Klebsiella pneumoniae*, *Pseudomonas aeruginosa, Yersinia ruckeri*, *Listeria monocytogenes*, *Staphylococcus lentus* and *Enterococcus faecalis*. Such an approach facilitates the systematic optimization of enzyme variants aimed at improving antibacterial performance.

Pre-ZooA_agal and ZooA_agal targets not only all evaluated *streptococci* species but also *Enterococcus faecalis,* a pathogen that accounts for up to 60% of the enterococcal mastitis cases^45^. While some endolysins possess a highly restricted host range, such as those targeting *S. suis*^12^, others, including PlyC, λSA2, B30, and the engineered chimeric lysin NC5, effectively cross-lyse multiple mastitis-causing species, namely *S. uberis*, *S. agalactiae*, and *S. dysgalactiae*^41,46^. Interestingly, reported recently homologous enzyme, SALys78925, differing only in 5 amino acids from pre-ZooA_agal, displayed similar specificity with the exception of *S. dysgalactiae* that was not affected by the enzyme treatment^47^.

### Efficacy in milk and serum

Turbidity reduction assays (TRAs) confirmed that pre-ZooA_agal and ZooA_agal exert rapid, dose-dependent lysis against susceptible strains, reducing the optical density of bacterial cultures by up to 70% and achieving a low minimum bactericidal concentration (MBC) of 0.5 μM. Both, pre-ZooA_agal and a mature enzyme - ZooA_agal, demonstrated high bactericidal efficacy, achieving a 3-to 4-log_10_ reduction (up to 99.99% bacterial cell eradication) from an initial cell density of 10^7^ CFU/ml. These results compare favorably with previously characterized streptococcal endolysins. While known streptococcal endolysins, such as PlySK1249 and PlySs9, typically achieve reductions of only 1.5 to 2.5 log_10_ within standard buffers^12^, and often require screening over 80,000 engineered candidates to identify a single variant with partial resilience in milk, the 3-to 4-log_10_ reduction reported herein for pre-ZooA_agal and ZooA_agal was successfully maintained within complex, physiologically relevant matrices.

It stands in stark contrast to previously characterized streptococcal endolysins, for which matrix-dependent loss of activity represents a major translational barrier^12,41^. Our finding that bacteriolytic efficacy was sustained in various matrix type (FBS, human serum, and UHT milk), demonstrates that these novel enzymes inherently possess the robust physiological features required for effective veterinary applications.

The sustained efficacy of pre-ZooA_agal and ZooA_agal in milk may stem from the unique fact that their lytic activity is not inhibited by the divalent metal ions inherently present in milk, such as Ca^2+^ and Mg^2+^. In stark contrast, several staphylolytic and streptococcolytic enzymes have been shown to be negatively affected by calcium and magnesium cations. Notably, Akesson et al. (2007)^18^, reported that the activity of the parental *Streptococcus equi*-derived zoocin A was markedly reduced in the presence of Ca^2+^ and Mg^2+^ions. The ability of pre-ZooA_agal and its mature derivative to maintain, and even enhance, their lytic performance in the presence of Ca^2+^ and Mg^2+^ represents a critical evolutionary and functional deviation from the zoocin A, explaining their superior and robust performance within the challenging milk and serum environments.

This stable performance positions our enzymes favourably not only relative to many wild-type endolysins described to date, but also places them within a comparable efficacy range to several engineered constructs developed specifically to overcome milk- and serum-associated inhibition.

### In vitro and in vivo safety profile

Safety is a paramount feature of novel antimicrobials. Here, *in vitro* MTT assays on NIH 3T3 fibroblasts and HaCaT keratinocytes confirmed the absence of cytotoxicity for both pre-ZooA_agal and ZooA_agal to mammalian cells. Interestingly, pre-ZooA_agal induced dose-dependent mortality in *G. mellonella* larvae (reaching 90 % at 4 µM), while the mature ZooA_agal, as expected, caused only a negligible increase in mortality (<10 %) relative to buffer-injected controls. Further evaluation of ZooA_agal toxicity using zebrafish embryos model demonstrated non-toxic and non-teratogenic properties of ZooA_agal, even when applied at very high (4 µM) concentration.

Ultimately, the absence of acute toxicity, embryotoxicity, or teratogenicity of ZooA_agal variant across both vertebrate models underscores the favourable safety profile typically attributed to PGHs ^34,44,48^. This safety is inherently tied to their highly specialized mechanism of action, which exclusively targets bacterial cell wall components entirely absent in eukaryotic cells, highlighting the potential of ZooA_agal as a secure enzybiotic candidate for controlling streptococcal infections in veterinary medicine.

### Ultrastructural insights into cell wall disruption

Electron microscopy provided direct ultrastructural evidence of the bacteriolytic mode of action of ZooA_agal against *S. agalactiae*. While untreated control cells retained intact cell walls and homogeneous cytoplasmic contents, exposure to ZooA_agal resulted in profound and progressive ultrastructural deterioration. SEM revealed extracellular accumulation of cellular debris consistent with cytoplasmic leakage, whereas TEM confirmed extensive cell wall damage, including visible ruptures, alongside the appearance of electron-lucent ghost cells with disorganized cytoplasmic contents and fragmented wall remnants. These observations are consistent with a mechanism of bacterial killing driven by direct enzymatic degradation of the peptidoglycan layer, leading to a loss of structural integrity, membrane disintegration, and ultimately cytoplasmic efflux.

Strikingly similar ultrastructural perturbations have been described for the zoocin A-treated *S. pyogenes* cells, including cytoplasmic disorganization and eventual complete cell wall rupture, concluded that bacterial killing occurs through direct enzymatic lysis^49^. The strong correspondence between those published findings and the ultrastructural alterations observed here for ZooA_agal, strongly supports the conclusion that the bacteriolytic mechanism is fully conserved between these two homologous enzymes.

### Translational perspectives

The potent, selective bacteriolytic activity and favourable safety profiles of pre-ZooA_agal and ZooA_agal underscore their potential as targeted enzybiotic candidate for veterinary applications. Their narrow spectrum of activity is particularly advantageous for controlling streptococcal pathogens such as *S. agalactiae*, *S. uberis* and *S. dysgalactiae*. This selectivity is of paramount relevance in the context of the bovine mammary microbiota, which is now recognized as a complex and functionally significant microbial community rather than a sterile environment. Bovine milk harbours a resident microbiota whose core microorganisms contribute to mammary health and milk quality, with origins traced both to the mammary gland itself and to entero- and rumen-mammary pathways^50^.

Disruption of this commensal community by broad-spectrum antimicrobials carries severe consequences by altering the overall microbial composition of bovine milk^51^. The targeted mode of action of pre-ZooA_agal and ZooA_agal, directed specifically against relevant pathogenic bacteria while sparing the commensal community, represents a distinct advantage over conventional antibiotic treatment. Such specificity minimizes collateral disruption of the beneficial mammary microbiota and mitigates the selective pressures that drive the emergence and persistence of antimicrobial resistance, aligning closely with One Health principles and current global efforts to reduce antibiotic dependency in dairy production.

Potential future applications of these variants include enzyme-based intramammary therapeutics for mastitis, topical formulations for wound infections, or incorporation into veterinary hygiene and biosecurity protocols. Future studies should evaluate their *in vivo* efficacy in mammalian infection models and further explore their performance in complex biological environments relevant to veterinary practice.

## Declarations

### Ethics approval and consent to participate

All experimental procedures involving animals were conducted in accordance with applicable national and institutional guidelines for the care and use of laboratory animals. Experiments using *D. rerio* embryos were performed in compliance with the regulations governing the use of early life-stage zebrafish embryos, which are not considered protected animals under European Directive 2010/63/EU prior to independent feeding. Experiments with *G. mellonella* larvae were conducted according to established laboratory practices for invertebrate models and did not require formal ethical approval.

### Consent for publication

Not applicable.

### Availability of data and materials

The datasets generated and/or analyzed during the current study are available from the corresponding author on reasonable request.

### Authors’ contributions

MN designed the study, developed the experimental methodology, performed protein expression, purification, and biochemical characterization experiments, conducted antimicrobial activity assays and data analysis, analyzed and interpreted the data, and drafted the manuscript. MKD conceived and designed the study and contributed to data interpretation. MK performed and described toxicity experiments. MG performed electron microscopy experiments and contributed to the interpretation of microscopy data. IS supervised the project, acquired funding, and contributed to manuscript writing and editing. All authors contributed to revising the manuscript critically for important intellectual content and approved the final version of the manuscript.

## Acknowledgements

The authors express their gratitude to Ryszard Strzalkowski (IMDiK PAN) for preparing and visualizing samples using scanning electron microscopy and Monika Matuszczyk for technical assistance and support during this study.

## References

1. Martins, L. et al. Association between antimicrobial use and antimicrobial resistance of Streptococcus uberis causing clinical mastitis. J. Dairy Sci. 104, 12030–12041 (2021).

2. Wente, N. et al. Associations between Streptococcus uberis strains from the animal environment and clinical bovine mastitis cases. J. Dairy Sci. 102, 9360–9369 (2019).

3. Dobrut, A. et al. Molecular and phenotypic identification of bacterial species isolated from cows with mastitis from three regions of Poland. BMC Vet. Res. 20, 193 (2024).

4. Brouwer, S. et al. Pathogenesis, epidemiology and control of Group A Streptococcus infection. Nat. Rev. Microbiol. 21, 431–447 (2023).

5. Alves-Barroco, C. et al. New Insights on Streptococcus dysgalactiae subsp. dysgalactiae Isolates. Front. Microbiol. 12, 686413 (2021).

6. Uruén, C., García, C., Fraile, L., Tommassen, J. & Arenas, J. How Streptococcus suis escapes antibiotic treatments. Vet. Res. 53, 91 (2022).

7. Gray, J. A. et al. Novel Biocontrol Methods for Listeria monocytogenes Biofilms in Food Production Facilities. Front. Microbiol. 9, 605 (2018).

8. Loessner, M. J. Bacteriophage endolysins — current state of research and applications. Curr. Opin. Microbiol. 8, 480–487 (2005).

9. Bustamante, N. et al. Deciphering how Cpl-7 cell wall-binding repeats recognize the bacterial peptidoglycan. Sci. Rep. 7, 16494 (2017).

10. Liu, H. et al. Therapeutic potential of bacteriophage endolysins for infections caused by Gram-positive bacteria. J. Biomed. Sci. 30, 29 (2023).

11. Georgakis, N. et al. The impact of metagenomic analysis on the discovery of novel endolysins. Appl. Microbiol. Biotechnol. 109, 126 (2025).

12. Vander Elst, N. Bacteriophage-derived endolysins as innovative antimicrobials against bovine mastitis-causing streptococci and staphylococci: a state-of-the-art review. Acta Vet. Scand. 66, 20 (2024).

13. Kranjec, C. et al. An antibiotic-free antimicrobial combination of bacteriocins and a peptidoglycan hydrolase: *in vitro* and *in vivo* assessment of its efficacy. Appl. Environ. Microbiol. 91, e02433–24 (2025).

14. Totté, J. et al. Targeted anti-staphylococcal therapy with endolysins in atopic dermatitis and the effect on steroid use, disease severity and the microbiome: study protocol for a randomized controlled trial (MAAS trial). Trials 18, 404 (2017).

15. De Wit, J. et al. Endolysin treatment against Staphylococcus aureus in adults with atopic dermatitis: A randomized controlled trial. J. Allergy Clin. Immunol. 144, 860–863 (2019).

16. ClinicalTrials.gov. First-in-Man Single-Dose and Multiple Dose Study to Evaluate the Safety, Tolerability and Efficacy of HY-133 (HY-133). (2024).

17. Simmonds, R. S., Simpson, W. J. & Tagg, J. R. Cloning and sequence analysis of zooA, a Streptococcus zooepidemicus gene encoding a bacteriocin-like inhibitory substance having a domain structure similar to that of lysostaphin. Gene 189, 255–261 (1997).

18. Akesson, M., Dufour, M., Sloan, G. L. & Simmonds, R. S. Targeting of streptococci by zoocin A. FEMS Microbiol. Lett. 270, 155–161 (2007).

19. Lai, A. C.-Y., Tran, S. & Simmonds, R. S. Functional characterization of domains found within a lytic enzyme produced by *Streptococcus equi* subsp. *zooepidemicus*. FEMS Microbiol. Lett. 215, 133–138 (2002).

20. Gargis, S. R. et al. Use of 4-Sulfophenyl Isothiocyanate Labeling and Mass Spectrometry To Determine the Site of Action of the Streptococcolytic Peptidoglycan Hydrolase Zoocin A. Appl. Environ. Microbiol. 75, 72–77 (2009).

21. Sabala, I. et al. Crystal structure of the antimicrobial peptidase lysostaphin from *Staphylococcus simulans*. FEBS J. 281, 4112–4122 (2014).

22. Odintsov, S. G., Sabala, I., Marcyjaniak, M. & Bochtler, M. Latent LytM at 1.3Å Resolution. J. Mol. Biol. 335, 775–785 (2004).

23. Chen, Y., Simmonds, R. S., Young, J. K. & Timkovich, R. Solution structure of the recombinant target recognition domain of zoocin A. Proteins Struct. Funct. Bioinforma. 81, 722–727 (2013).

24. Kim, H., Lee, H. G., Kwon, I. & Seo, J. Characterization of Endolysin LyJH307 with Antimicrobial Activity against Streptococcus bovis. Animals 10, 963 (2020).

25. Ning, H. et al. Gut virome derived bacteriophage endolysin Lys572 can effectively control Streptococcus suis in refrigerated and thawed pork. Int. J. Food Microbiol. 452, 111707 (2026).

26. Vidal Amaral, J. R., et al. Bacteriocin Producing Streptococcus agalactiae Strains Isolated from Bovine Mastitis in Brazil. Microorganisms 10, 588 (2022).

27. Gargis, S. R. et al. Zif, the Zoocin A Immunity Factor, Is a FemABX-Like Immunity Protein with a Novel Mode of Action. Appl. Environ. Microbiol. 75, 6205–6210 (2009).

28. Gargis, A. S., O’Rourke, A.-L. D., Sloan, G. L. & Simmonds, R. S. Prevalence and Acquisition of the Genes for Zoocin A and Zoocin A Resistance in Streptococcus equi subsp. zooepidemicus. J. Mol. Evol. 68, 498–505 (2009).

29. Mirdita, M. et al. ColabFold: making protein folding accessible to all. Nat. Methods 19, 679–682 (2022).

30. Jumper, J. et al. Highly accurate protein structure prediction with AlphaFold. Nature 596, 583–589 (2021).

31. Mirdita, M. et al. Uniclust databases of clustered and deeply annotated protein sequences and alignments. Nucleic Acids Res. 45, D170–D176 (2017).

32. Li, M. Z. & Elledge, S. J. Harnessing homologous recombination in vitro to generate recombinant DNA via SLIC. Nat. Methods 4, 251–256 (2007).

33. Hartmann, M. et al. Damage of the Bacterial Cell Envelope by Antimicrobial Peptides Gramicidin S and PGLa as Revealed by Transmission and Scanning Electron Microscopy. Antimicrob. Agents Chemother. 54, 3132–3142 (2010).

34. Kaus-Drobek, M. et al. From discovery to potential application: engineering a novel M23 peptidase to combat Listeria monocytogenes. Sci. Rep. 15, 15628 (2025).

35. E3 medium (for zebrafish embryos). Cold Spring Harb. Protoc. 2011, pdb.rec66449 (2011).

36. Jones, P. et al. InterProScan 5: genome-scale protein function classification. Bioinformatics 30, 1236–1240 (2014).

37. Zhang, M.-J. et al. Insights into the Influence of Signal Peptide on the Enzymatic Properties of Alginate Lyase AlyI1 with Removal Effect on Pseudomonas aeruginosa Biofilm. Mar. Drugs 20, 753 (2022).

38. Schober, I. et al. Bac *Dive* in 2025: the core database for prokaryotic strain data. Nucleic Acids Res. 53, D748–D756 (2025).

39. Moradi, A. et al. Trends in Antimicrobial Resistance of Major Mastitis-Causing Pathogens: A Nine-Year Study. *Vet*. Med. Sci. 11, e70417 (2025).

40. Kabelitz, T., Aubry, E., Van Vorst, K., Amon, T. & Fulde, M. The Role of Streptococcus spp. in Bovine Mastitis. Microorganisms 9, 1497 (2021).

41. Vander Elst, N., et al. Development of engineered endolysins with in vitro intracellular activity against streptococcal bovine mastitis-causing pathogens. Microb. Biotechnol. 16, 2367–2386 (2023).

42. Shen, Y. et al. A bacteriophage endolysin that eliminates intracellular streptococci. eLife 5, e13152 (2016).

43. Wong, K. Y. et al. Endolysins against Streptococci as an antibiotic alternative. Front. Microbiol. 13, 935145 (2022).

44. Mitkowski, P. et al. Enhancing stability and safety of chimeric peptidoglycan hydrolases by linker engineering. Appl. Microbiol. Biotechnol. 110, 21 (2026).

45. Różańska, H., Lewtak-Piłat, A., Kubajka, M. & Weiner, M. Occurrence of enterococci in mastitic cow’s milk and their antimicrobial resistance. *J*. Vet. Res. 63, 93–97 (2019).

46. Schmelcher, M., Powell, A. M., Camp, M. J., Pohl, C. S. & Donovan, D. M. Synergistic streptococcal phage λSA2 and B30 endolysins kill streptococci in cow milk and in a mouse model of mastitis. Appl. Microbiol. Biotechnol. 99, 8475–8486 (2015).

47. Yoon, S., Moon, J., Kim, H. & Seo, J. Development and characterization of endolysin SALys78925 with antimicrobial activity against Streptococcus agalactiae. J. Anim. Sci. Technol. https://doi.org/10.5187/jast.2025.e42 (2025) doi:10.5187/jast.2025.e42.

48. Małecki, P. H. et al. Novel endopeptidase overcoming lysostaphin resistance. Sci. Rep. https://doi.org/10.1038/s41598-026-43937-3 (2026) doi:10.1038/s41598-026-43937-3.

49. Simmonds, R. S., Pearson, L., Kennedy, R. C. & Tagg, J. R. Mode of action of a lysostaphin-like bacteriolytic agent produced by Streptococcus zooepidemicus 4881. Appl. Environ. Microbiol. 62, 4536–4541 (1996).

50. Guo, W. et al. Bovine milk microbiota: Key players, origins, and potential contributions to early-life gut development. J. Adv. Res. 59, 49–64 (2024).

51. Winther, A. R., Perrin, A., Nordraak, A. O. O., Kjos, M. & Porcellato, D. An in vitro evaluation of the effect of antimicrobial treatment on bovine mammary microbiota. Sci. Rep. 14, 18333 (2024).

