## Supplementary material for "Characterization and evaluation of the anti-streptococcal activity of a novel zoocin A-like endopeptidase for potential veterinary applications"

**Supplementary Table S1**. Bacterial strains used in this study, their source and PG characterization. **DSMZ-** German Collection of Microorganisms and Cell Cultures; **CCM-** Czech Collection of Microorganisms; **ATCC-** American Type Culture Collection; **NCTC-** National Collection of Type Cultures; **PCM-** Polish Collection of Microorganisms.

| Bacterial strain and source | Source | Cross-bridge in peptidoglycan (PG) | PG chemotype | |
|---|---|---|---|---|
| | | | Schleifer & Kandler (1972)[1] | Schumann (2011)[2] |
| ***Gram-negative strains*** | | | | |
| *Escherichia coli* DSM 1103; clinical isolate (Seattle, 1946) | DSMZ | meso-DAP–D-Ala | A1γ | A31 |
| *Klebsiella pneumoniae* DSM 789; cow's milk | DSMZ | | | |
| *Pseudomonas aeruginosa* DSM 939; animal room water bottle | DSMZ | | | |
| *Yersinia ruckeri* CCM 4620; rainbow trout with red-mouth disease | CCM | | | |
| ***Gram-positive strains*** | | | | |
| *Bacillus subtilis* DSM 10; type strain Marburg | DSMZ | meso-DAP–D-Ala | A1γ | A31 |
| *Corynebacterium striatum* DSM 20668; unknown | DSMZ | | | |
| *Listeria monocytogenes* DSM 19094; poultry | DSMZ | | | |
| *Lactococcus lactis* CCM 1877; type strain | CCM | L-Lys–D-Asp | A4α | A11.31 |
| *Enterococcus faecium* DSM 2146; cheese | DSMZ | | | |
| *Staphylococcus aureus* NCTC 8325-4; derivative of NCTC 8325, originally isolated from a human corneal ulcer | NCTC | L-Lys–Gly(5) | A3α | A11.3 |
| *Staphylococcus lentus* PCM 2441; goat udder | PCM | L-Lys–Gly(4–5) | A3α | A11.2 |
| *Staphylococcus epidermidis* ATCC 12228; FDA strain PCI 1200 | ATCC | L-Lys–Gly$_{2-4}$–L-Ser$_{1-2}$–Gly | A3α | A11.3 |

| Bacterial strain and source | Source | Cross-bridge in peptidoglycan (PG) | PG chemotype | |
|---|---|---|---|---|
| | | | Schleifer & Kandler (1972)[1] | Schumann (2011)[2] |
| *Staphylococcus simulans* CCM 3583; unknown | CCM | | | |
| *Micrococcus luteus* ATCC 10240; air | ATCC | L-Lys peptide subunit | A2 | A11.pep |
| *Enterococcus faecalis* DSM 20376; raw milk | DSMZ | L-Lys–L-$Ala_2$ | A3α | A11.5 |
| *Streptococcus agalactiae* DSM 6784; bovine udder infection | DSMZ | | | |
| *Streptococcus dysgalactiae* DSM 20662; cow | DSMZ | | | |
| *Streptococcus pyogenes* CCM 7418; pharynx of a child | CCM | | | |
| *Streptococcus equi subsp. zooepidemicus* DSM 20272; isolated from animals and their surroundings | DSMZ | | | |
| *Streptococcus salivarius* DSM 20560; blood, from a patient with acute articular rheumatism | DSMZ | L-Lys–L-$Ala_2$ | A3α | A11.5 |
| *Streptococcus uberis* DSM 20569; unknown | DSMZ | | | |
| *Streptococcus canis* DSM 20715; bovine mastitis | CCM | L-Lys–L-Thr–Gly | A3α | A11.24 |

**Supplementary Table S2.** Primers used for genes cloning

| Primer Name | Primer sequence (5'-3') |
|---|---|
| 1FL_IF | CAAACCGGTGGATCCATGAACAAGTGGCTGGTTAAAGC |
| 1FL_IR | GTGGTGGTGCTCGAGTTATTATTTATACGGCAGTTCCTG |
| 1FL_WF | CCAGCCACTTGTTCATGGATCCACCGGTTTGTTCCTGG |
| 1FL_WR | CAGGAACTGCCGTATAAATAACTCGAGCACCACCACCAC |
| 1Cat_TRD IF | CAAACCGGTGGATCCGTTCGCCCGATTGATAATGGC |
| 1CatTRD_VF | CAATCGGGCGAACGGATCCACCGGTTTGTTCCTG |

**Supplementary Figure S1**. Structural analysis of pre-ZooA-agal protein. Cartoon representation of AlphaFold model of pre-ZooA_agal; the colour corresponds to the colouring of the domains in Fig. 1: grey for signal peptide (SP), orange for M23 and green for TRD. Binding groove is indicated with dotted circle.

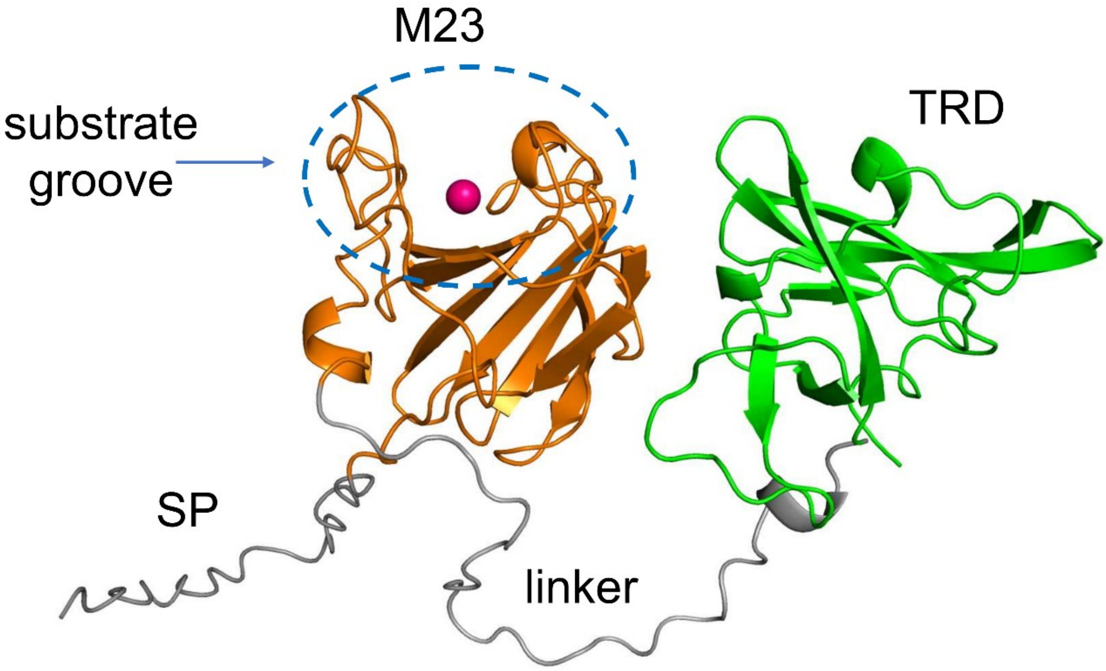

**Supplementary Figure S2.** Antibacterial activity of pre-ZooA_agal against *S. agalactiae* in biologically relevant matrices. The bacteria were incubated with 5µM enzyme in the presence of human serum, fetal bovine serum (FBS) or UHT milk and afterwards samples were serially diluted and spotted on TSA plates. A) Activity measured after 1h incubation in serum and milk diluted to 25% or 12.5% final concentration. B) Activity measured after 24h incubation in 100% or 50% in serum or milk at room temperature.

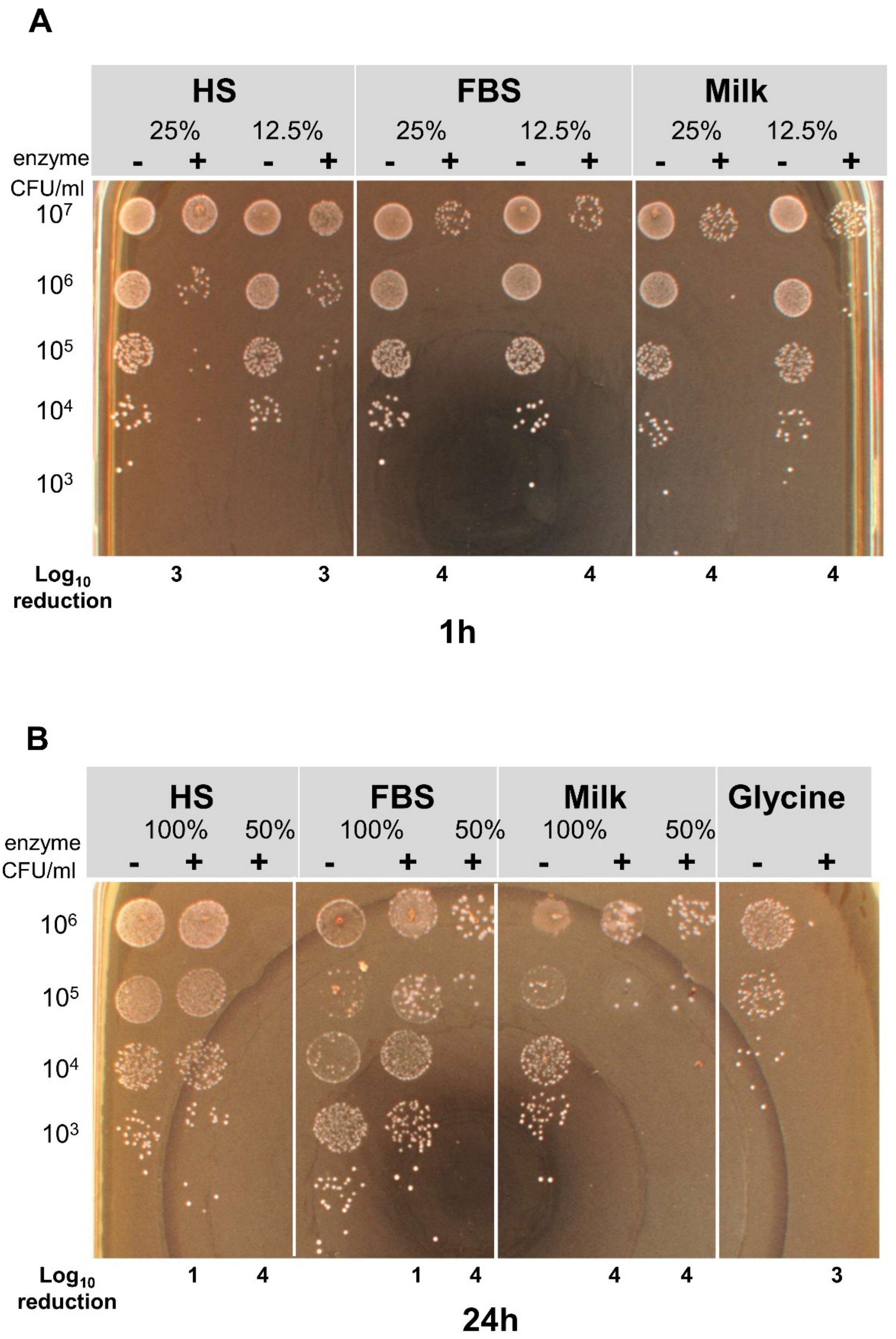

**Supplementary Figure S3.** Determination of the minimum bactericidal concentration (MBC) as the lowest enzyme concentration where no growth of bacteria was detected in spot killing assay against. *S. agalactiae, S. uberis and S. dysgalactiae* $10^5$ CFU/ml cells were treated with ZooA_agal at concentration between 0.0011 to 1 µM in a 50 mM glycine (pH 8.0) for 24 h at RT. Bacteria strains were plated alone as a control.

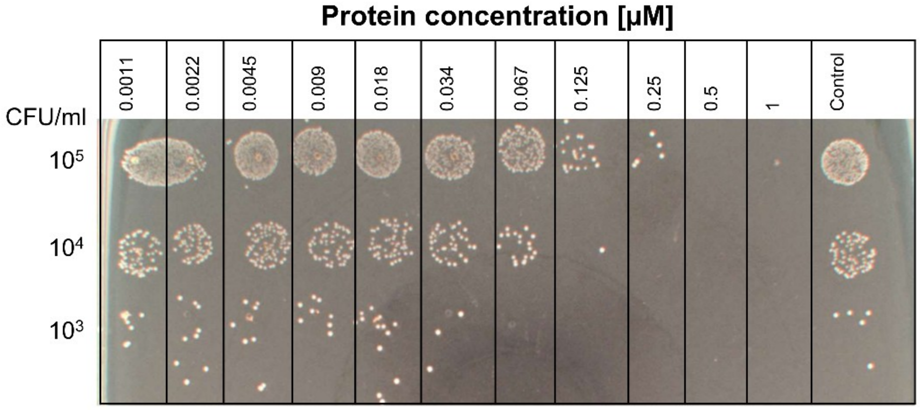


*S.agalactiae*

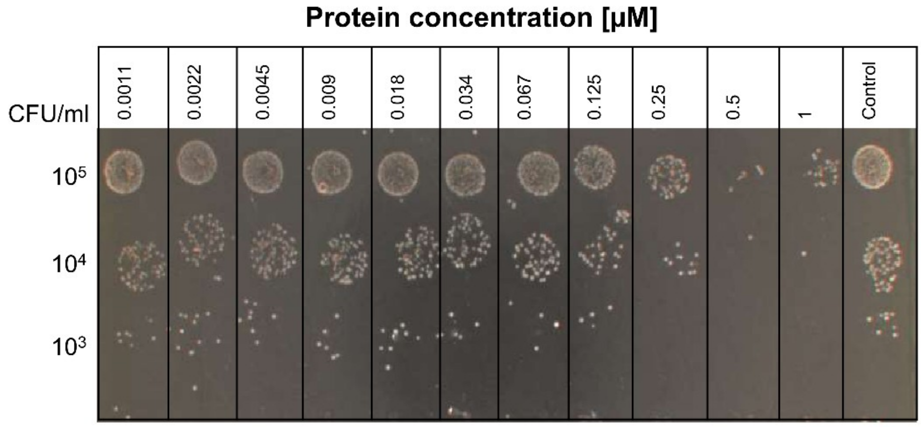


*S.uberis*

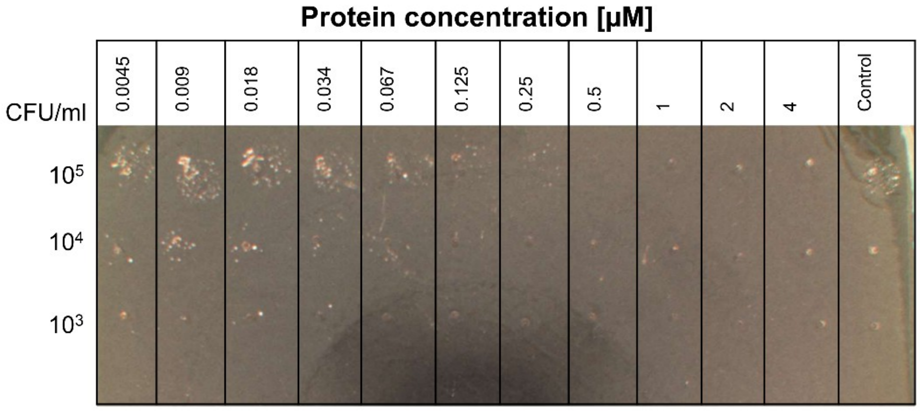


*S.dysgalactiae*

**References:**

1. Schleifer, K. H. & Kandler, O. Peptidoglycan types of bacterial cell walls and their taxonomic implications. *Bacteriol. Rev.* **36**, 407–477 (1972).

2. Schumann, P. Peptidoglycan Structure. in *Methods in Microbiology* vol. 38 101–129 (Elsevier, 2011).
